# Long-term efficacy of adoptive cell therapy is determined by host CD8^+^ T cells and undermined by lymphodepleting preconditioning

**DOI:** 10.1101/2024.05.16.594554

**Authors:** Diego Figueroa, Juan Pablo Vega, Andrés Hernández-Oliveras, Felipe Gálvez-Cancino, Felipe Ardiles, Felipe Flores, Sofía Hidalgo, Ximena López, Hugo Gonzalez, Fabiola Osorio, Vincenzo Borgna, Alvaro Lladser

## Abstract

Adoptive T cell therapy (ACT) has demonstrated remarkable efficacy in treating hematological cancers. However, its efficacy against solid tumors remains limited and the emergence of cancer cells that lose expression of targeted antigens often promotes resistance to ACT. Importantly, the mechanisms underlying effective and durable ACT-mediated tumor control are incompletely understood. Here, we show that adoptive transfer of TCR-transgenic CD8^+^ T cells eliminates established murine melanoma tumors, with concomitant accumulation of tumor-infiltrating CD8^+^ T cells exhibiting both progenitor-exhausted and terminally-differentiated phenotypes. Interestingly, host CD8^+^ T cells contributed to ACT-mediated elimination of primary tumors and rejected ACT-resistant melanoma cells lacking the targeted antigen. Mechanistically, ACT induced TNF-α- and cross-presenting dendritic cell-dependent tumor accumulation of endogenous CD8^+^ T cells and effective tumor elimination. Importantly, although lymphodepleting preconditioning enhanced ACT-mediated tumor elimination, it abrogated host antitumor immunity and protection against ACT-resistant melanoma cells. Enrichment of transcriptional signatures associated with TNF-α signaling, cross-presenting dendritic cells and tumor-specific CD8^+^ T cells in human melanoma tumors correlated with favorable responses to ACT and increased survival. Our findings reveal that long-term efficacy of ACT is determined by the interplay between transferred and endogenous CD8^+^ T cells and is undermined by lymphodepleting preconditioning, which ultimately favors ACT resistance.

## INTRODUCTION

Antitumor immunity is largely mediated by tumor-specific CD8^+^ T cells, which are activated in the lymph nodes by migratory conventional type 1 dendritic cells (cDC1) cross-presenting tumor antigen-derived peptides onto major histocompatibility complex (MHC) class I molecules (1–3). CD8^+^ T cells that recognize tumor antigen/MHC-I complexes through their T cell receptor (TCR), proliferate, and differentiate into cytotoxic CD8^+^ T cells that then migrate to tumors, where they recognize target cancer cells and kill them by releasing cytotoxic molecules, such as perforin, granzymes and granulysin (4). In addition, they also secrete effector cytokines, such as IFN-γ and TNF-α, which promote antigen presentation on target cells and activate myeloid cells, including macrophages and DCs (5,6). In the tumor microenvironment, chronic TCR stimulation induces a dysfunctional differentiation program in tumor-specific CD8^+^ T cells, known as T cell exhaustion, which is characterized by the expression of the transcription factor TOX and multiple inhibitory receptors, such as PD-1 (7,8). Exhausted CD8^+^ T cells comprise at least two functionally distinct populations: progenitor-exhausted (Tpex) and terminally-differentiated (Tdif) cells (9,10). The Tpex subset expresses the transcription factor TCF-1, has self-renewal potential, and upon antigen-mediated TCR activation, give rise to Tdif cells. The latter up-regulate cytotoxic molecules and the inhibitory receptor TIM-3, but lack TCF-1 and self-renewal potential (9). In addition, both Tpex and Tdif populations acquire a tissue-resident transcriptional program that allows them to adapt and establish in different microenvironment (10). Accumulating evidence indicates that an effective antitumor immunity relies on the concerted action of both Tpex (PD1^+^TOX^+^TCF-1^+^GzmB^-^ TIM3^-^) and Tdif (PD1^high^TOX^+^TCF-1^-^GzmB^+^TIM3^+^) cells (11–13). Interestingly, better clinical outcomes depend on the ability of Tpex cells to lead the tumor accumulation of Tdif cells (14). All these studies provide evidence supporting that the Tpex pool is important for long-term establishment in tumors, whereas Tdif cells are critical for exerting cytotoxic antitumor activity.

Adoptive cell therapy (ACT), which consists of the infusion of autologous tumor-specific T cells expanded *ex vivo*, has emerged as a novel treatment for hematological cancers and has shown promise in the treatment of solid tumors (15). ACT using tumor-infiltrating lymphocytes (TILs) has been shown to induce therapeutic activity with curative potential in patients with melanoma and other solid tumors (16,17). However, the implementation of reproducible TIL therapies remains challenging due to the low frequency, dysfunctional state, and limited proliferation of tumor-reactive T cells (18). On the other hand, ACT strategies using peripheral blood-derived T cells genetically engineered to express tumor-specific TCR or chimeric antigen receptors (CAR) have overcome these limitations and have demonstrated potent antitumor responses (19,20). CAR-T cell immunotherapies have shown remarkable success in patients with different hematological cancers, including subtypes of leukemia, lymphoma, and myeloma (21,22) Currently, six different CAR-T cell immunotherapies are approved by the F.D.A. for hematologic cancers but none for solid tumors (23). TCR-based ACTs have been tested in clinical studies for almost two decades, showing safety and promising antitumor responses in patients with metastatic melanoma and other solid tumors (24). Despite significant advances in the field, most cancer patients with solid tumors fail to respond to ACT or develop resistance through different mechanisms. An important factor contributing to ACT efficacy is the persistence of transferred T cells *in vivo* (25). Consequently, strategies aimed at improving T cell persistence have been extensively used. Non-myeloablative lymphodepletion as a preconditioning regimen, such as chemotherapy and/or radiation, is an integral part of currently used clinical protocols. Such regimens favor the engraftment of transferred T cells, probably by attenuating the competition between infused and endogenous T cells for cytokines and a niche (26). In addition, the appearance of mutant tumor cells that lack the expression of targeted antigens is a major mechanism of resistance to ACT (27–29). Antigen loss can occur due to several mechanisms ranging from genetic to post-translational alterations (30,31)(32). In this scenario, the immune system acts as a selective pressure that leads to the selection of tumor cell clones with decreased immunogenicity, e.g. lacking the target antigen (33,34).

Emerging evidence indicates that stimulating the host immune system can help to overcome these mechanisms of ACT resistance (35). In fact, combining ACT with oncolytic viruses, Toll-like receptor agonists and/or DC activators, that expand endogenous tumor-specific CD8^+^ T cells can provide long-lasting tumor protection and overcome the appearance of antigen-loss escape variants (36,37). However, whether transferred T cells can intrinsically engage the participation of endogenous antitumor T cells to promote effective elimination of primary tumors and provide long-lasting immunity against ACT-resistant tumor cells remains unknown. Here, we show that ACT with TCR-transgenic CD8^+^ T cells engage the participation of endogenous CD8^+^ T cells exhibiting tumor-specific phenotypes to effectively reject established syngeneic melanoma tumors in mice. Moreover, ACT-promoted host CD8^+^ T cell immunity protected against rechallenge with ACT-resistant melanoma cells lacking the targeted antigen. Mechanistically, ACT induced TNF-α-and cross-presenting dendritic cell-dependent tumor accumulation of endogenous CD8^+^ T cells and effective tumor elimination. Interestingly, preexistence of transcriptional signatures associated with TNF-α signaling, cross-presenting DCs and tumor-specific CD8^+^ T cells in human melanoma tumors correlated with favorable responses to ACT and increased survival. Our findings reveal a TNF-α-and cDC1-dependent interplay between transferred and endogenous CD8^+^ T cells that determines the efficacy of ACT to eradicate solid tumors and provide long-term antitumor immunity, which is undermined by lymphodepleting preconditioning, ultimately favoring acquired resistance to ACT.

## RESULTS

### Effective ACT promotes tumor accumulation of endogenous CD8^+^ T cells exhibiting Tpex and Tdif phenotypes

We established an ACT model consisting of intravenous administration of *in vitro*-activated OTI TCR-transgenic CD8^+^ T cells to mice bearing B16F10-OTI melanoma tumors (∼50-200 mm^3^) engineered to express the ovalbumin-derived epitope OTI (SIINFEKL) (38) (Fig. 1a). In this setting, tumors were efficiently eliminated within approximately ten days, providing long-term survival (Fig. 1b, c). To understand the mechanisms underlying the effective antitumor immunity observed in this ACT model, we analyzed the tumor cell infiltrates by flow cytometry three days after adoptive transfer, which correspond to the regression phase of tumor growth. We analyzed the frequencies and phenotypes of endogenous (CD45.1^-^) and transferred (CD45.1^+^) CD8^+^ T cells (Fig. 1d). We observed that ACT increased total CD45^+^ hematopoietic cells in tumors (Fig. 1d). Within the CD45^+^ cell population, we observed a prominent accumulation of endogenous CD8^+^ T cells, as compared to untreated controls (Fig. 1d, e). In particular, we observed an accumulation of PD-1^+^GzmB^-^ and PD-1^+^GzmB^+^ subpopulations (Fig. 1d, e), which correspond to progenitor exhausted (Tpex) and terminally differentiated (Tdif) phenotypes of tumor-reactive CD8^+^ T cells, respectively (11). In addition, Tpex cells expressed TCF1 and TOX1, whereas Tdif cells expressed TOX-1 and TIM3 (Fig. 1f). Transferred OTI cells displayed an effector phenotype characterized by intermediate levels of PD-1 and granzyme B, as well as low levels of TOX-1 and TCF-1 (Fig. 1f), as compared to endogenous subsets (39,40). These results show that effective ACT promotes accumulation of tumor-resident CD8^+^ T cells, including both Tpex (PD1^+^, GzmB^-^, TCF1^+^, TOX^+^ and TIM3^-^) and Tdif subsets (PD1^+^, GzmB^+^, TCF1^-^, TOX^+^ and TIM3^+^), which have been shown to mediate antitumor immunity and predict favorable response to immunotherapy (12). In contrast, a suboptimal ACT protocol that initially reduces tumor growth but is not able to eliminate tumors, did not increase the frequencies of endogenous CD8^+^ T cells (Supplementary Fig. 1). These observations led us to hypothesize that ACT engages the participation of endogenous tumor-specific CD8^+^ T cells to mediate effective tumor control.

**Figure 1.**
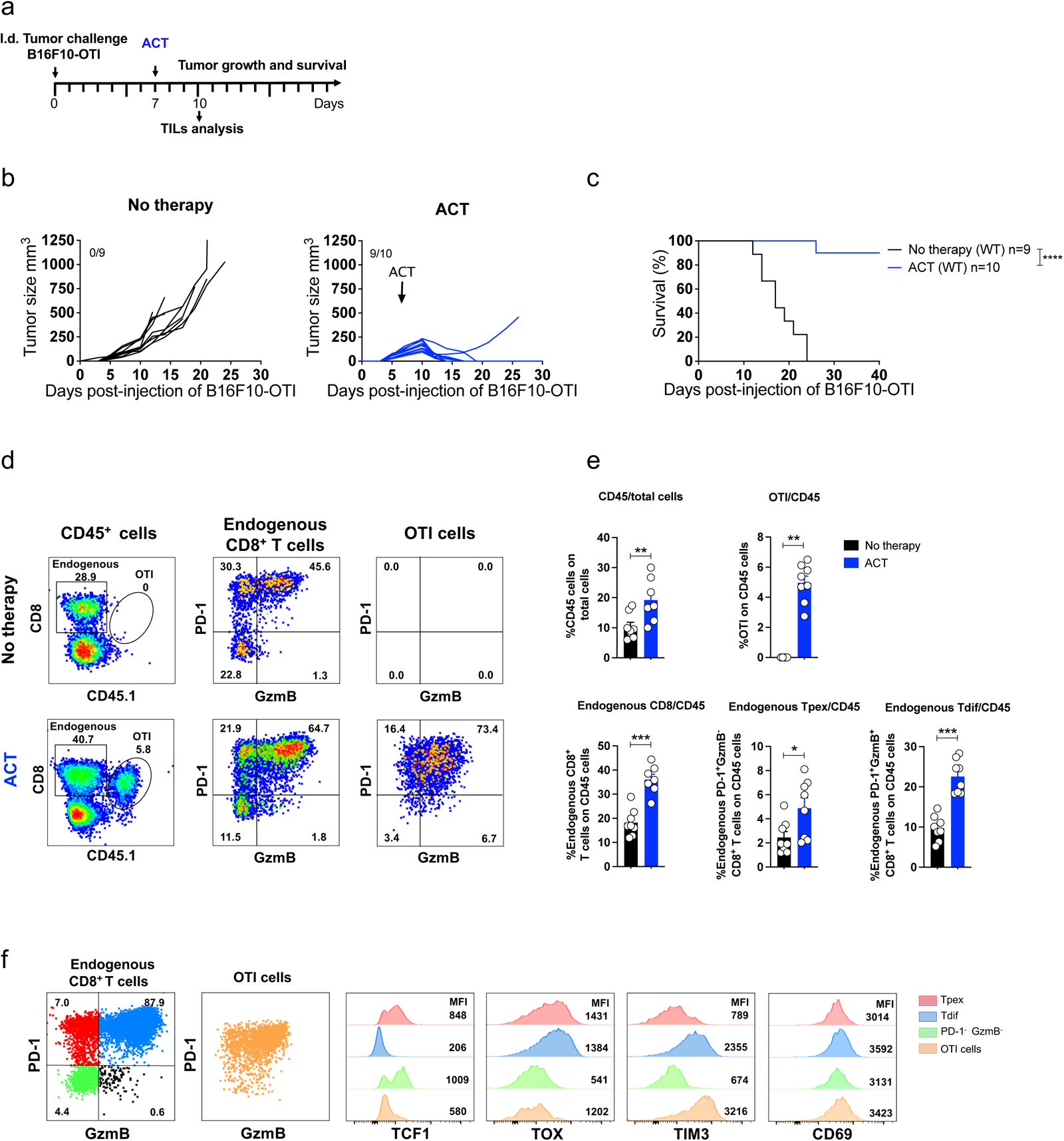
ACT induces rejection of melanoma tumors and tumor accumulation of endogenous CD8^+^ T cells with Tpex and Tdif phenotypes. C57BL/6 mice bearing B16F10-OTI tumors received i.v. transfer of 1x10^6^ *in vitro* activated OTI CD8^+^ T cells (ACT). Untreated mice (No therapy) were used as controls. **a** Experimental scheme. **b-c** Individual tumor growth (**b**) and Kaplan-Meier (**c**) curves for each group: No therapy (black curves), and ACT (blue curves). **d-f** Tumor infiltrates were analyzed by flow cytometry three days after ACT. **d** Left panels: Representative dot plots displaying the frequencies of endogenous (square, CD45.1^-^) and transferred (ellipse, OTI CD45.1^+^) CD8^+^ T cells in live CD45^+^ cells. Middle panels: Representative dot plots displaying the expression of PD-1 and granzyme B in endogenous CD8^+^ T cells and the frequencies of the different subpopulations defined in each quadrant: PD-1^+^ GzmB^-^ (Tpex), PD-1^+^ GzmB^-^ (Tdif), PD-1^-^ GzmB^-^ and PD-1^-^ GzmB^+^. Right panels: Representative dot plots displaying the expression of PD-1 and granzyme B in transferred OTI CD8^+^ T cells. **e** Quantifications of CD45^+^ cells as a percentage of total live cells, as well as transferred OTI, total endogenous, Tpex (PD-1^+^ GzmB^-^) and Tdif (PD-1^+^ GzmB^+^) CD8^+^ T cells as a percentage of CD45^+^ cells. **f** Representative dot plots displaying PD-1 and GzmB expression in endogenous CD8^+^ T cells and OTI cells. Histograms show the expression of TCF-1, TOX and TIM3 in the different subpopulations defined in each quadrant: Endogenous PD-1^+^ GzmB^-^ (Tpex, red), Endogenous PD-1^+^ GzmB^-^ (Tdif, blue), Endogenous PD-1^-^ GzmB^-^ (green) and transferred OTI cells (orange). (**c**) Kaplan-Meier curve shows survival of each condition of two independent experiment n=9-10. *p ˂0.05, **p ˂0.01, ***p ≤ 0.001, and ****p ≤ 0.0001 by Log-rank Mantel-Cox test. (**e**) Pooled data from two independent experiments, n = 8 per group. Bars are the mean ± SEM. *p<0.05, **p< 0.01, ***p< 0.001 by Mann-Whitney unpaired.

### Endogenous CD8^+^ T cells contribute to ACT-mediated elimination of primary tumors and reject ACT-resistant melanoma cells

We investigated the contribution of host immune system to ACT-mediated tumor control using complementary approaches. First, we tested the ability of ACT to control tumor growth in RAGKO mice, which lack mature T and B cells (Supplementary Fig. 2a). In these mice, tumors grew faster than in immunocompetent wild-type mice, and ACT failed to control tumor growth (Supplementary Fig 2b, c), arguing that endogenous T and/or B cells are required for the antitumor effects of ACT. To confirm the contribution of endogenous T cells in immunocompetent mice, we tested ACT in mice treated with FTY720 (Fig. 2a), which prevented circulation (data not shown) and tumor accumulation of endogenous T cells without affecting tumor infiltration of transferred CD8^+^ T cells (Fig. 2b, c) (41). We observed that ACT was unable to control tumor growth in FTY720-treated mice (Fig. 2d, e), indicating that endogenous T cells play a key role in the efficacy of ACT. To evaluate whether host CD8^+^ T cells are directly involved in supporting the antitumor effects of ACT, mice were treated with anti-CD8 depleting antibodies prior to the adoptive transfer (Fig. 3a), which efficiently eliminate CD8^+^ T cells (Fig. 3b, c). Interestingly, ACT was not able to control tumor growth efficiently in mice depleted of endogenous CD8^+^ T cells, and the protection in terms of survival was completely abolished (Fig. 3d, e). In contrast, CD4^+^ T cell depletion did not impair the ability of ACT to reject tumors (data not shown). To assess the ability of host antitumor immunity to reject ACT-resistant melanoma cells, mice that had eliminated B16F10-OTI melanoma tumors were rechallenged in the opposite flank with wild-type B16F10 cells (Fig. 3f). As hypothesized, most of these mice were protected (Fig. 3g, h), and this protection was abolished when CD8^+^ T cells were depleted prior to rechallenge (Fig. 3g, h), indicating that effective ACT promotes long-term CD8^+^ T cell-mediated host immunity against other melanoma antigens, a phenomenon known as antigen spreading. This response is relevant for controlling tumor cells that lose the expression of the targeted antigen, a well-documented mechanism of ACT resistance leading to disease progression (42). Taken together, these results highlight the ability of ACT to engage the participation of endogenous CD8^+^ T cells to collectively provide potent and broad long-lasting antitumor immunity.

**Figure 2.**
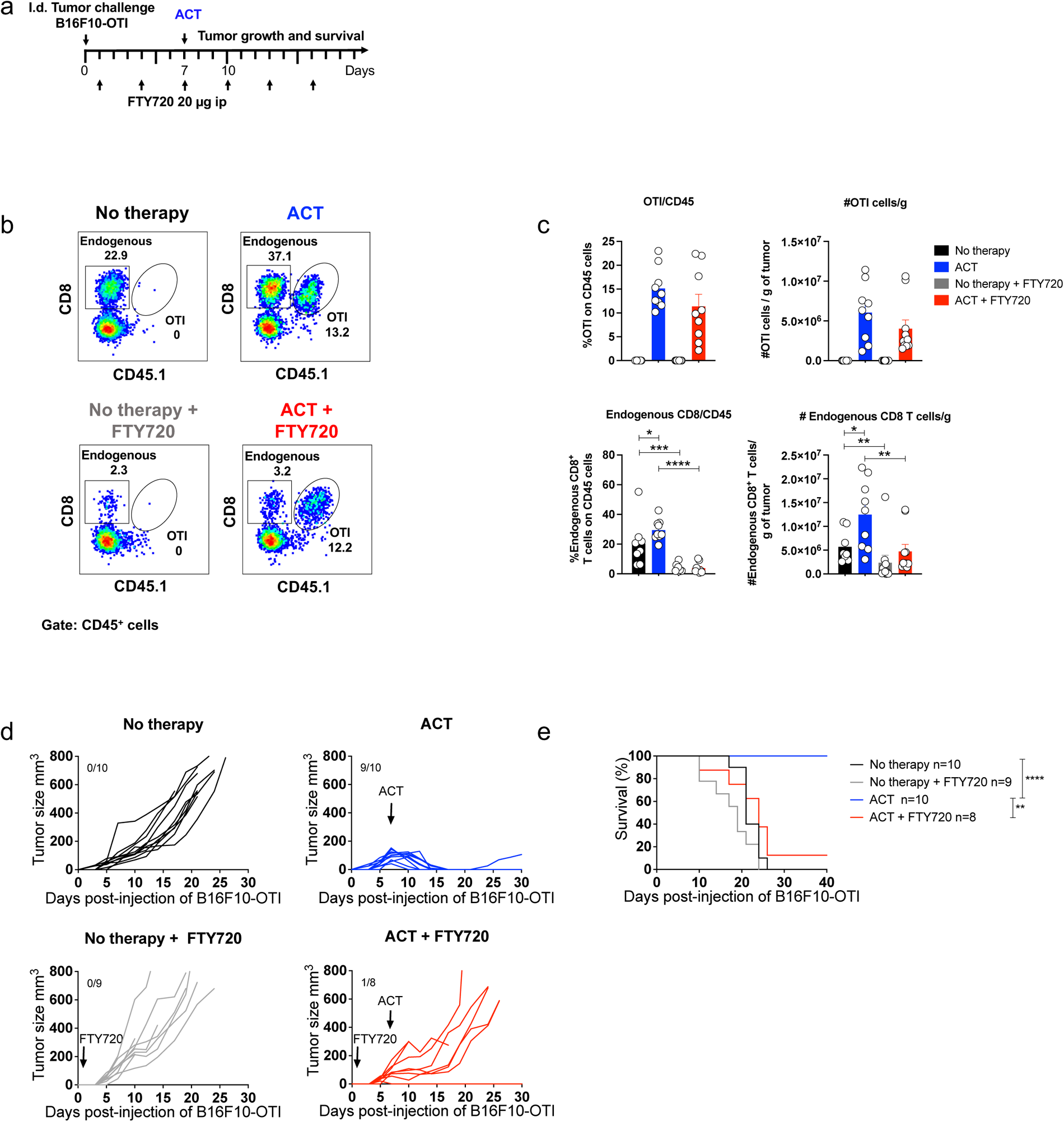
Endogenous T cells are required for ACT-induced tumor protection. WT C57BL/6 mice bearing B16F10-OTI tumors received i.p. injections of FTY720 every three days starting one day after the tumor challenge, then where i.v. transfer with 1x10^6^ *in vitro* activated OTI CD8^+^ T cells (ACT). **a** Experimental scheme**. b-c** Tumor infiltrating lymphocytes were analyzed by flow cytometry three days after ACT. **b** Representative dot plots displaying the frequencies of endogenous (square, CD45.1^-^) and transferred (ellipse, OTI CD45.1^+^) CD8^+^ T cells in live CD45^+^ cells for each group: No therapy, ACT, No therapy + FTY720 and ACT + FTY720. **c** Relative (on CD45^+^ cells) and absolute quantifications of transferred OTI (upper panel) and endogenous (lower panel) CD8^+^ T cells for each group: No therapy (black bars), ACT (blue bars), No therapy + FTY720 (grey bars), ACT + FTY720 (red bars). **d-e** Individual tumor growth (**d**) and Kaplan-Meier (**e**) curves for each group: No therapy (black curves), ACT (blue curves), No therapy + FTY720 (gray curves), ACT + FTY720 (red curves). (**c**) Pooled data from two independent experiments, n=9-10 per group. Bars are the mean ± SEM. **p ˂0.05, **p ˂0.01, ***p ≤ 0.001, and ****p ≤ 0.0001 by Mann-Whitney unpaired test. (**e**) Results from two independent experiment n=8-10. *p ˂0.05, **p ˂0.01, ***p ≤ 0.001, and ****p ≤ 0.0001 by Log-rank Mantel-Cox test.

**Figure 3.**
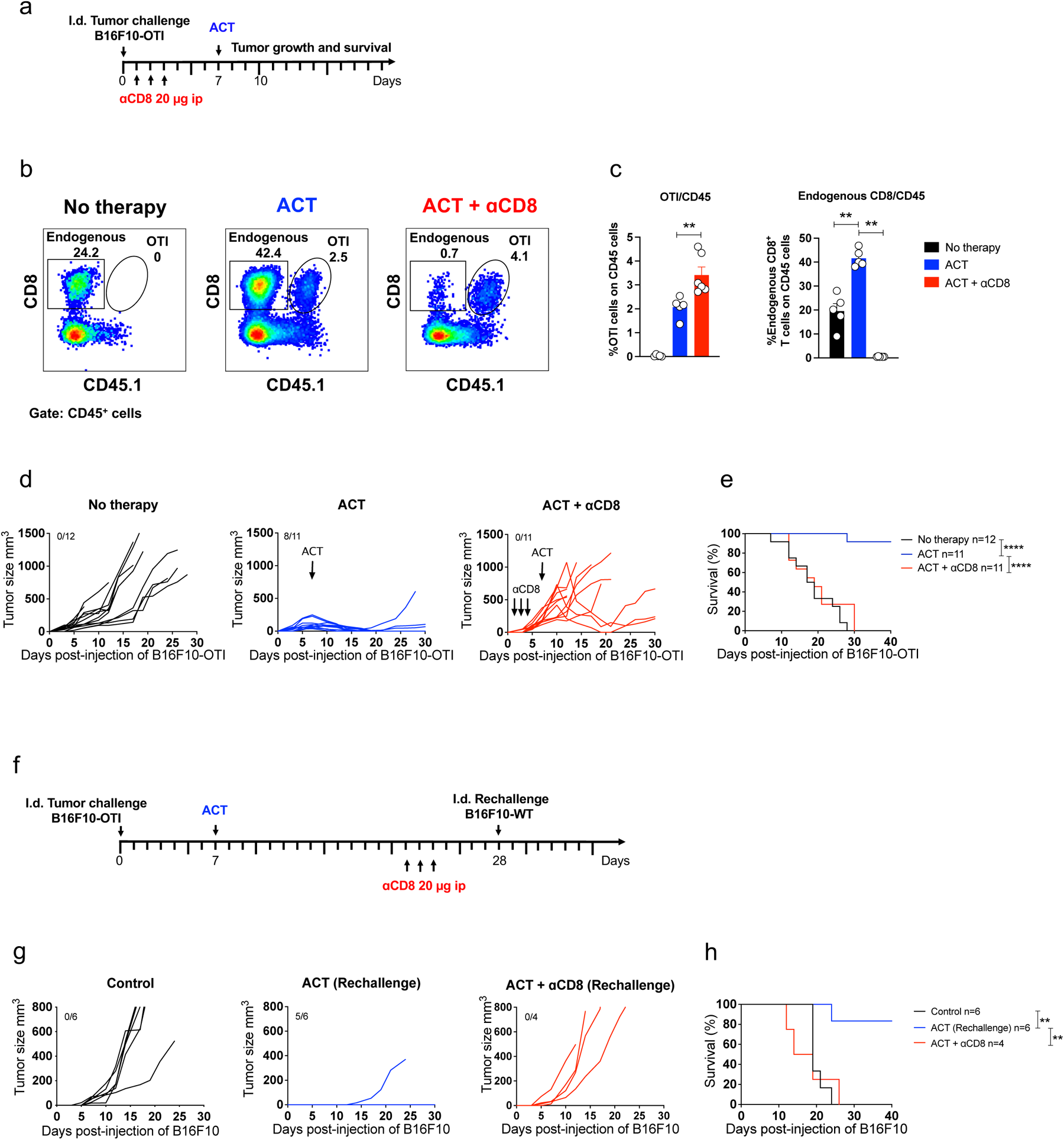
Endogenous CD8^+^ T cells are essential for ACT efficacy. C57BL/6 mice bearing B16F10-OTI tumors received i.v. transfer of 1x10^6^ *in vitro* activated OTI CD8^+^ T cells (ACT). Untreated mice (No therapy) were used as controls. Some groups of mice received three i.p. daily doses of anti-CD8 antibodies starting one day after tumor challenge. **a** Experimental timeline. **b-c** Tumor infiltrates were analyzed by flow cytometry three days after ACT. **b** Representative dot plots displaying the frequencies of endogenous (square, CD45.1^-^) and transferred (ellipse, OTI CD45.1^+^) CD8^+^ T cells in live CD45^+^ cells for each group: No therapy, ACT and ACT + anti-CD8. **c** Quantification of transferred OTI (left panel) and endogenous (right panel) CD8^+^ T cells as the percentage of CD45^+^ cells for each group: No therapy (black bars), ACT (blue bars), ACT + anti-CD8 (red bars). **d-e** Individual tumor growth (**d**) and Kaplan-Meier (**e**) curves for each group: No therapy (black curves), ACT (blue curves), and ACT + anti-CD8 (red curves). Mice that rejected B16F10-OTI tumors after ACT received or did not receive three i.p. daily doses of anti-CD8 and 5 days later were i.d. rechallenged with 1x10^6^ of B16F10 WT cell lines in the opposite flank. **f** Experimental timeline **g-h** Individual tumor growth (**g**) and Kaplan-Meier curves (**h**) for tumor re-challenged groups: Control (black curves) ACT (blue curves) and ACT + anti-CD8 (red curves). (**c**) one experiments, n=5-6 per group. Bars are the mean ± SEM. **p ˂0.05, **p ˂0.01, ***p ≤ 0.001, and ****p ≤ 0.0001 by Mann-Whitney unpaired test. (**d-e**) Data from three independent experiments n=11-12 and (**h**) from one experiment n=4-6. *p ˂0.05, **p ˂0.01, ***p ≤ 0.001, and ****p ≤ 0.0001 by Log-rank Mantel-Cox test.

### ACT induces TNF-α- and cross-presenting dendritic cell-dependent tumor accumulation of endogenous CD8^+^ T cells and effective tumor elimination

To investigate the mechanism underlying the interplay between transferred and endogenous CD8^+^ T cells, we examined the involvement of TNF-α, an effector cytokine that is known to promote innate and adaptive antitumor immune responses (43). For this purpose, anti-TNF-α blocking antibody was administered starting one day before ACT (Fig. 4a). As expected, TNF-α blockade prevented ACT-induced tumor accumulation of endogenous CD8^+^ T cells exhibiting Tpex and Tdif phenotypes (Fig. 4b) and significantly reduced the elimination of primary tumors and mouse survival (Fig. 4c). These results suggest that TNF-α may promote tumor accumulation of endogenous CD8^+^ T cells via activation of cross-presenting cDC1, which present tumor-derived antigens to tumor-specific CD8^+^ T cells in draining lymph nodes (44,45). Hence, we analyzed DCs in tumors and draining lymph nodes (Fig. 4a; Supplementary Fig. 3) and observed that upon ACT, tumor-infiltrating cDC1 were decreased (Fig. 4d) but exhibited higher maturation levels (frequencies of PDL1^high^, CD86^high^ and CD86^high^PDL1^high^ cDC1) (Fig. 4e). These effects were impeded in mice treated with anti-TNF-α antibody (Fig. 4d, e), suggesting that ACT promotes maturation and migration to draining lymph nodes of cDC1 in a TNF-α-dependent manner. As expected, ACT induced a TNF-α-dependent increase in the frequency of migratory cDC1 in draining lymph nodes (Fig. 4f), particularly those exhibiting a higher maturation phenotype CD86^high^PDL1^high^ (Fig. 4g). To address the participation of cDC1 in the expansion of endogenous CD8^+^ T cells, we used the Langerin-DTR mice that express the human diphtheria toxin receptor (DTR) and the enhanced green fluorescent protein (EGFP) genes under the control of langerin (CD207) promoter (46). In this transgenic mouse model, DTR and EGFP are expressed at different levels among different DC subsets (Supplementary Fig. 4a), allowing the depletion of tissue-migratory and lymph node-resident cDC1s, as well as Langerhans cells after diphtheria toxin (DTx) administration (Supplementary Fig. 4b, c). In these mice, depletion of tumor cDC1 was not efficient, which likely reflects their relatively low DTR expression (Supplementary Fig. 4a) and ensures the trafficking of transferred effector T cells to tumors (47). In Lang-DTR mice that started receiving DTx administrations one day prior to adoptive transfer (Fig. 5a), we observed that OTI CD8^+^ T cells efficiently infiltrated tumors (Fig. 5b), however, ACT was unable to promote tumor accumulation of endogenous CD8^+^ T cells displaying Tpex and Tdif phenotypes, as compared to wild-type mice (Fig. 5b). Accordingly, ACT induced a transient antitumor effect in Lang-DTR mice but the majority of tumors progressed (Fig. 5c) in contrast to complete protection observed in wild-type mice (Fig. 5d). Taken together, these results demonstrate that effective ACT induces a TNF-α-and cDC1-dependent accumulation of intratumoral endogenous CD8^+^ T cells and tumor elimination.

**Figure 4.**
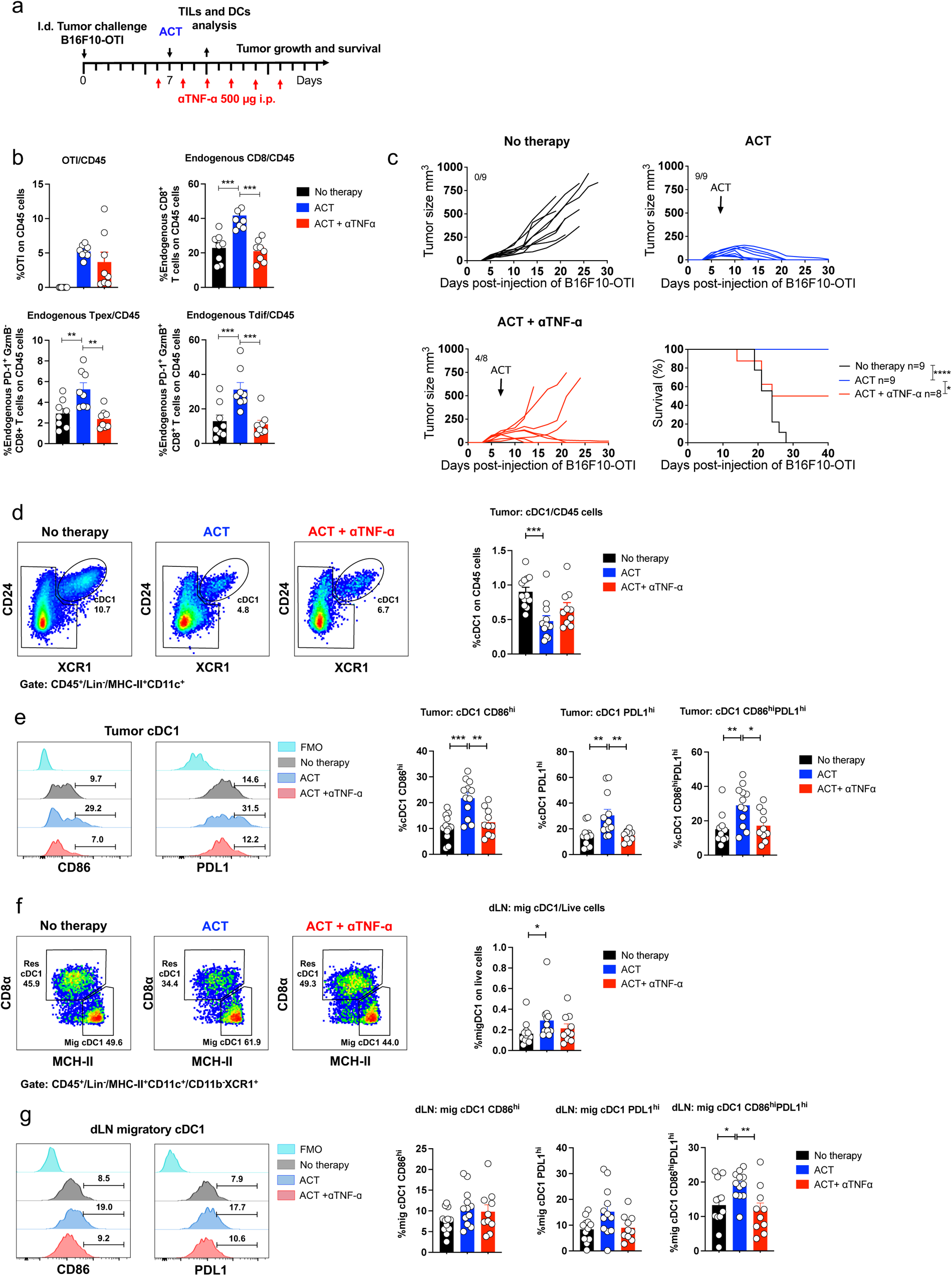
ACT induces TNF-α-dependent tumor accumulation of endogenous CD8^+^ T cells and cDC1 maturation and migration to draining lymph nodes. C57BL/6 mice bearing B16F10-OTI tumors received i.v. transfer of 1x10^6^ *in vitro* activated OTI CD8^+^ T cells (ACT). Untreated mice (No therapy) were used as controls. One group of ACT-treated mice received anti-TNF-α antibody i.p. every other day, starting one day before ACT. **a** Experimental design. **b** Quantification of tumor-infiltrating transferred OTI CD8^+^ T cells, endogenous CD8^+^ T cells, Tpex and Tdif cells on CD45^+^ cells of all condition analyzed by flow cytometry three days after ACT. **c** Tumor growth and Kaplan-Meier curves for each group: No therapy (black curves), ACT (blue curves) and ACT + anti-TNF-α (Red curves). **d-g** Tumors (**d, e**) and draining lymph nodes (**f, g**) were analyzed by flow cytometry three days after ACT for each group: No therapy, ACT, ACT + anti-TNF-α. **d** Representative dot plots and their respective quantification displaying the frequencies of tumor-infiltrating cDC1 (ellipse, CD24^+^XCR1^+^) within CD45^+^/Lin^-^/MHCII^+^CD11c^+^ cells. **e** Histograms showing CD86 and PDL1 expression in tumor-infiltrating cDC1, and frequency quantifications of CD86^hi^ PDL1^hi^ cDC1 for each group: No therapy (black bars), ACT (blue bars), ACT + anti-TNF-α (red bars). **f** Representative dot plots and their respective quantification displaying the frequencies of resident (CD8α^+^ and MHC class II^+^) and migratory (CD8α^-^ and MHC class II^high^) cDC1 within CD45^+^/Lin^-^/MHCII^+^CD11c^+^/CD11b^-^XCR1^+^ in tumor-draining lymph nodes. **g** Histograms showing the expression of CD86 and PDL1 in migratory cDC1, and frequency quantifications of CD86^hi^PDL1^hi^ cDC1. (**b**) Pooled data of two independent experiments, n=8 per group. Bars are the mean ± SEM. **p ˂0.05, **p ˂0.01, ***p ≤ 0.001, and ****p ≤ 0.0001 by Mann-Whitney unpaired test. (**c**) Kaplan-Meier curve shows survival of each condition of two independent experiment n=8-9. *p ˂0.05, **p ˂0.01, ***p ≤ 0.001, and ****p ≤ 0.0001 by Log-rank Mantel-Cox test. (**d-g**) Pooled data from three independent experiments, n = 10-12 per group. Bars are the mean ± SEM. **p ˂0.05, **p ˂0.01, ***p ≤ 0.001, and ****p ≤ 0.0001 by Mann-Whitney unpaired test.

**Figure 5.**
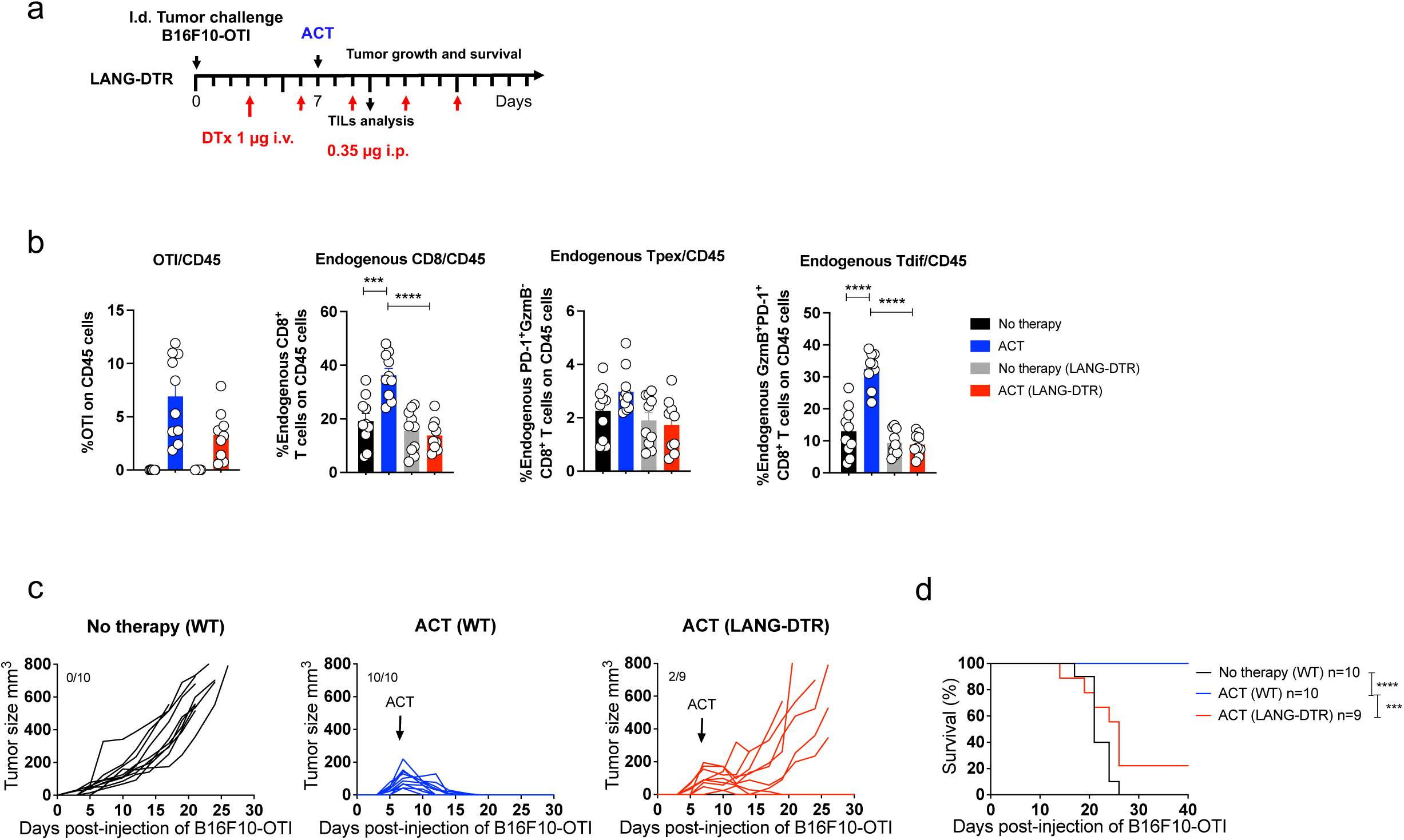
Tumor accumulation of endogenous CD8^+^ T cells and tumor elimination induced by ACT require cDC1. C57BL/6 and Langerin-DTR mice bearing B16F10-OTI tumors received i.v. transfer of 1x10^6^ *in vitro* activated OTI CD8^+^ T cells (ACT). Untreated mice (No therapy) were used as controls. Langerin-DTR mice received diphtheria toxin (DTx) every three days starting four days before the ACT. **a** Experimental timeline. **b** Quantifications of OTI, total endogenous, Tpex (PD-1^+^ GzmB^-^) and Tdif (PD-1^+^ GzmB^+^) CD8^+^ T cells as a percentage of CD45^+^ cells in tumors analyzed by flow cytometry three days after ACT for each group: No therapy (C57BL/6 WT mice: black bars; Lang-DTR mice: grey bars) and ACT (C57BL/6 WT mice: blue bars; Lang-DTR mice: red bars). **c-d** Individual tumor growth (**c**) and Kaplan-Meier (**d**) curves for each group: No therapy (C57BL/6 WT mice: black curves) and ACT (C57BL/6 WT mice: blue curves; Lang-DTR mice: red curves). (**b**) Pooled data from two independent experiments, n = 9-10 per group. Bars are the mean ± SEM. **p ˂0.05, **p ˂0.01, ***p ≤ 0.001, and ****p ≤ 0.0001 by Mann-Whitney unpaired test. (**d**) Kaplan-Meier curve shows survival from two independent experiment n=9-10. *p ˂0.05, **p ˂0.01, ***p ≤ 0.001, and ****p ≤ 0.0001 by Log-rank Mantel-Cox test.

### Lymphodepleting preconditioning enhances ACT-mediated tumor elimination, but abrogates long-term host antitumor immunity

Next, we assessed the significance of the interplay between transferred and endogenous CD8^+^ T cells in a clinically relevant setting. We studied the effects of lymphodepleting preconditioning, which is a component part of ACT protocols in the clinic, in our mouse ACT model. To this end, we administered two doses of cyclophosphamide (Cy) in consecutive days prior to ACT, which approximates clinical practice and has been widely used in the literature (48) (Fig. 6a). As expected, we observed that Cy induced a marked depletion of immune CD45^+^ cells (data not shown), including endogenous CD8^+^ T cells (Fig. 6b, c). Despite this, lymphodepleting preconditioning promoted the expansion of transferred OTI CD8^+^ T cells, resulting in efficient elimination of primary tumors, leading to complete responses in most mice tested (Fig. 6b-e). Furthermore, using a suboptimal ACT protocol, Cy also led to expansion of transferred cells and promoted tumor rejection in nearly all mice (Fig. 6b-e), indicating that lymphodepleting preconditioning contributes to effective control of primary tumors in these settings. To evaluate the impact of lymphodepleting preconditioning in the long-term protection against ACT-resistant melanoma cells, mice were rechallenged with B16F10 cells. Despite initial tumor control, mice treated with Cy and either optimal or suboptimal ACT protocol were unable to reject the rechallenge with B16F10 cells (Fig. 6f, g), indicating that lymphodepleting preconditioning has a detrimental effect on host antitumor immunity. These findings demonstrate that ACT promotes host antitumor immunity, which is severely compromised by lymphodepleting preconditioning regimen

**Figure 6.**
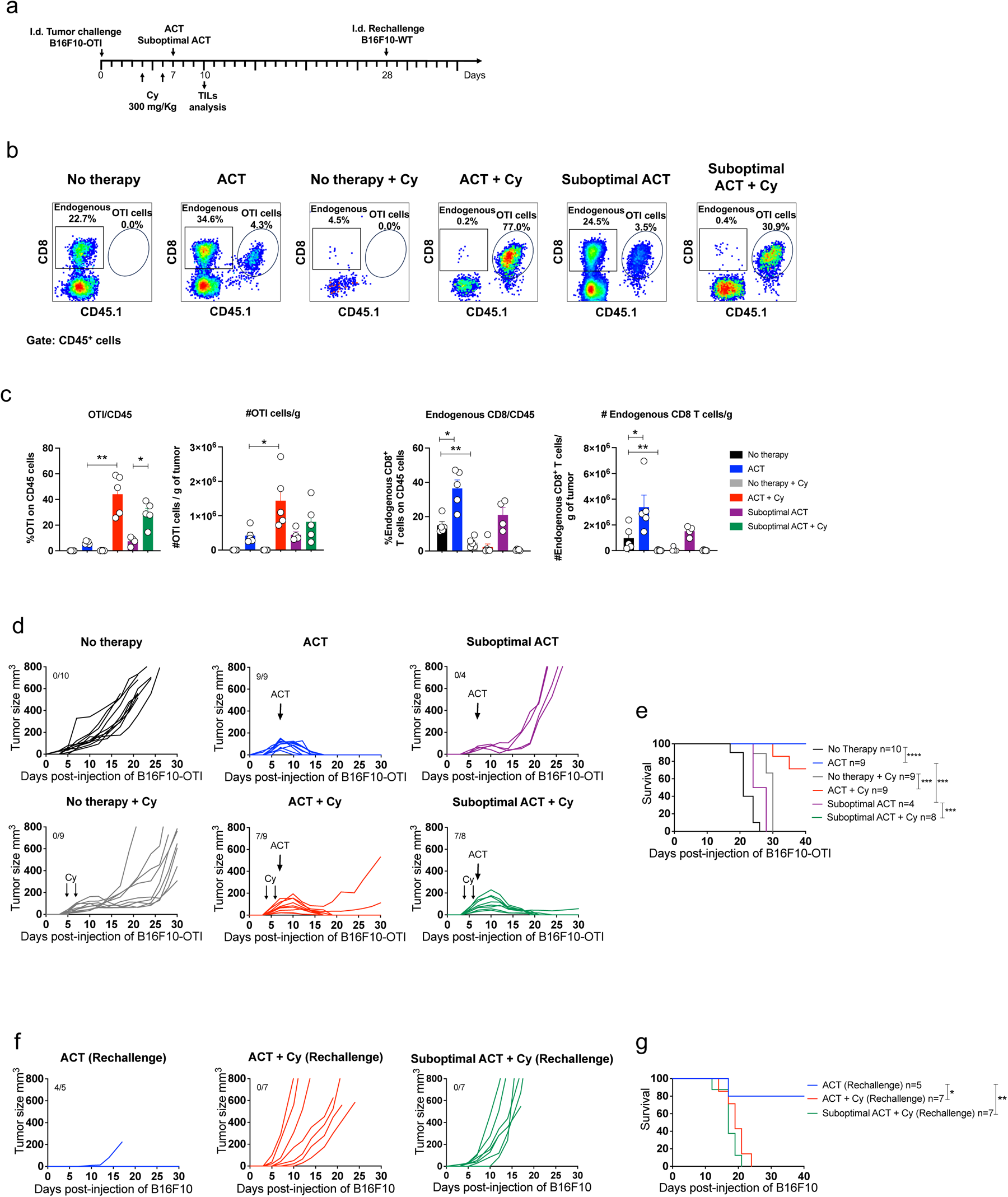
Lymphodepleting preconditioning enhances ACT-mediated tumor elimination, but abrogates long-term host antitumor immunity. C57BL/6 mice were intradermally challenged with 1x10^6^ B16F10-OTI, then mice received two doses of 300 mg/kg cyclophosphamide (Cy) on days 4 and 6 after tumor challenge as a preconditioning lymphodepleting regimen. One day later, mice received i.v. transfer of 1x10^6^ or 0.5x10^6^ *in vitro* activated OTI CD8^+^ T cells as optimal and suboptimal ACT models, respectively. Subsequently, all mice that rejected tumors after ACT were rechallenged with 1x10^6^ B16F10-WT in the opposite flank. Tumor growth and survival were analyzed every other day. **a** Experimental timeline. **b-c** Representative dot plots displaying the frequencies of endogenous (square, CD45.1^-^) and transferred (ellipse, OTI CD45.1^+^) CD8^+^ T cells in live CD45^+^ cells for each group: No therapy, ACT, No therapy + Cy, ACT + Cy, Suboptimal ACT and Suboptimal ACT + Cy (**c**) Relative (on CD45^+^ cells) and absolute quantification of transferred OTI and endogenous CD8^+^ T cells for each group shown in b. **d-e** Individual tumor growth (**d**) and Kaplan-Meier (**e**) curves for each group: No therapy (black curves), ACT (blue curves) and suboptimal ACT (purple curves) in non-lymphodepleted mice, and No therapy (gray curves), ACT (red curves) and suboptimal ACT (green curves) in lymphodepleted mice with 300 mg/kg of Cy. **f-g** Individual tumor growth (**f**) and Kaplan-Meier (**g**) curves of B16F10-WT rechallenged mice for each group: ACT (blue curves) in non-lymphodepleted, and ACT (red curves) and suboptimal ACT (green curves) in lymphodepleted mice with 300 mg/kg of Cy. Bars are the mean ± SEM. **p ˂0.05, **p ˂0.01, ***p ≤ 0.001, and ****p ≤ 0.0001 by Mann-Whitney unpaired test. (**e**) Kaplan-Meier curve showing the survival from two independent experiments, n=4-10 and (**g**) from one experiment, n=5-7. p ˂ 0.05, **p ˂ 0.01, ***p ≤ 0.001, and ****p ≤ 0.0001 by log-rank Mantel-Cox test

### cDC1, TNF-α signaling, Tpex and Tdif gene signatures positively associate with favorable clinical response to ACT and overall survival in human melanoma

Next, to investigate whether these findings might be relevant in humans, we analyzed publicly available datasets of melanoma patients receiving ACT for which tumor bulk RNAseq and clinical response (RECIST) data were available (49). Consistent with our findings in mice, the cDC1 gene signature was enriched in tumors of patients showing favorable clinical responses to ACT, including complete (CR) and partial (PR) responses, whereas it was down-regulated in patients with progressive (PD) and stable (SD) disease (Fig. 7a). In contrast, cDC2 and cDC3 signatures were enriched only in PR and not in CR patients. Non-specific signatures for active DC (acDC) or immature DC (immDC) showed no significant enrichment in PR or CR patients. These results show that among the different DC signatures, cDC1 is specifically associated with complete clinical response to ACT. The TNF-α -signaling gene signature was strongly associated with PR but weakly associated with CR, probably reflecting a far more complex network of signaling pathways involved in protective antitumor responses. Interestingly, Tpex and particularly Tdif gene signatures were enriched in patients who responded favorably to ACT, association that was stronger for CR patients (Fig. 7a). Also, we analyzed an immune gene signature upregulated in melanoma patients with immunotherapy resistance (ImmuneRes) (50), which was enriched in non-responder patients and decreased in responder patients. Finally, we investigated whether cDC1, TNF-α signaling, Tdif and Tpex gene signatures were associated with better survival in a larger cohort of patients with cutaneous skin melanoma (SKCM) from The Cancer Genome Atlas (TCGA) database. Higher expression levels of all these signatures were associated with longer survival. In contrast, high expression of the immune resistance signature showed a negative association with patient survival (Fig. 7b). These results support our findings in mouse models and suggest that cDC1, TNF-α signaling, Tpex and Tdif cells in tumors are indeed critical for the efficacy of ACT and the survival of melanoma patients.

**Figure 7.**
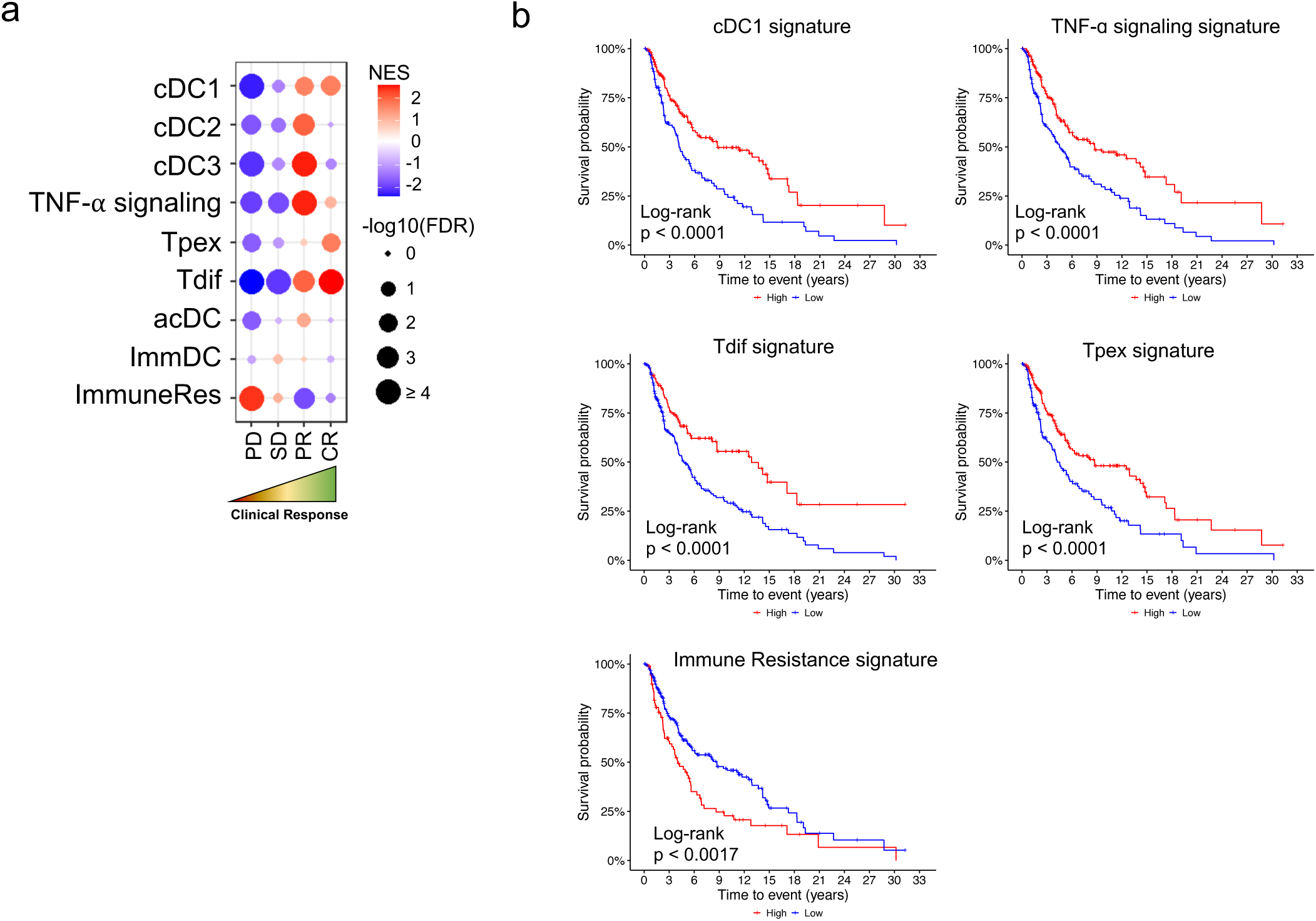
cDC1, TNF-α signaling, Tpex and Tdif gene signatures are positively associated with clinical responses to ACT and overall survival in melanoma patients. To explore the association between gene signatures and melanoma patients’ outcome, we analyzed gene expression data from patients under ACT, and from TCGA-SKCM cohort. **a** GSEA analysis in patients under ACT. Patients are divided according to RECIST clinical response. PD: progressive disease (n = 5); SD: stable disease (n = 10); PR: partial response (n = 5); CR: complete response (n = 5). To obtain the enrichment score, each RECIST response was compared against the others. NES: normalized enrichment score. An FDR < 0.05 (-log_10_ > 1.3) was considered as statistically significant. **b** Kaplan – Meier plots showing the overall survival analysis of TCGA-SKCM patients (n = 323) divided according to the high (red) and low (blue) expression levels of the indicated signature. p values were obtained with Log – rank test. p < 0.05 was considered as statistically significant.

## DISCUSSION

The reasons for the relatively poor efficacy and development of resistance of ACT in solid tumors are not yet clear (51). Here, we show that effective ACT engage the participation of endogenous CD8^+^ T cells to achieve potent and durable antitumor immunity. Using mouse models, we observed that adoptively transferred TCR-transgenic CTLs promoted a TNF-α- and cDC1-dependent expansion of host CD8^+^ T cells exhibiting Tpex and Tdif phenotypes. Furthermore, we demonstrated that endogenous CD8^+^ T cells are required for efficient elimination of primary melanoma tumors and rejection of ACT-resistant melanoma cells lacking the expression of the targeted antigen, which represent a major mechanism of ACT resistance is the emergence of antigen-loss mutants that escape immune recognition by ACT leading to cancer progression (27,42,52). Our findings also highlight the dual role of lymphodepleting preconditioning regimens which are used in the majority of ACT clinical protocols, including radiotherapy and chemotherapies such as cyclophosphamide and fludarabine (53). On the one hand, they can promote primary tumor elimination by promoting the expansion of transferred cells, as such regimens eliminate endogenous lymphocytes, reducing competition for niche and cytokines (54). On the other hand, lymphodepleting conditioning depletes endogenous tumor-reactive CD8^+^ T cells and other immune cells that support antitumor immunity, including cDC1, thereby compromising long-term host antitumor immunity, which is critical to combat ACT-resistant cancer cells. Our study uncovers a novel mechanism through which ACT mediates effective and durable tumor control and may underlie a major mechanism of acquired resistance to ACT in the clinic. Also, this study also may help to design optimized ACT protocols with improved efficacy for solid tumors.

Our work highlights the importance of the host immune system in the efficacy of ACT, which is emerging as a novel concept in the field. It was recently shown that optimal CAR-T cell activity relies on host IFN-γ-mediated signaling in a syngeneic mouse model of Burkitt-like lymphoma (55). Particularly, it was observed that CD4^+^ CAR-T cells promoted endogenous tumor-reactive CD8^+^ T cell responses, although their contribution to tumor elimination was not addressed (55). In contrast to this study, we provide evidence that TNF-α signaling plays a critical role in this ACT-host immunity crosstalk, which represents a novel mechanism. In line with this, a recent study showed that TNF-α produced by transferred CD8^+^ T cells was associated with better clinical responses to TCR-based ACT in metastatic melanoma patients (56). We also showed that TNF-α promoted ACT-induced maturation and migration to lymph nodes of cDC1, as well as tumor accumulation of endogenous Tpex and Tdif CD8^+^ T cells, resulting in enhanced tumor protection. This is consistent with the ability of CD8^+^ T cell-produced TNF-α to promote DC maturation in the context of viral infections (5) and with the role of migratory cDC1s carrying tumor-derived antigens to prime tumor-specific CD8^+^ T cells in draining lymph nodes (3). Indeed, we show that migratory and/or lymph node-resident cDC1s were needed for tumor accumulation of endogenous Tpex and Tdif CD8^+^ T cells and ACT efficacy, which is consistent with studies reporting that cDC1s maintain tumor-specific Tpex cells in tumor-draining lymph nodes, which then migrate to the tumor, where they differentiate into Tdif cells to sustain effective antitumor immunity (57–59). Our observations in mice align with analyses in RNAseq datasets from melanoma patients, where we found that TNF-α signaling, cDC1, Tpex and Tdif gene signatures in tumors positively correlate with therapeutic responses to ACT and overall survival. Taken together, our results support a key role of TNF-α and cDC1 in mediating the interplay between transferred and endogenous CD8^+^ T cells to achieve long-lasting protective immunity against solid tumors.

This study offers new insights into the crosstalk between adoptively transferred CD8^+^ T cells and the host immune system, which can be harnessed to improve the efficacy of ACTs against solid tumors. Consistent with this, some groups have combined ACT with strategies that activate and/or expand DCs, such as oncolytic viruses, TLR agonists, and DC activators (including FLT3L and CD40L). These strategies have been shown to broaden host antitumor T cell immunity and protect against antigen-loss escape variants (36,37,60). In contrast to these studies, we unveiled the intrinsic ability of transferred CD8^+^ T cells to mediate TNF-α-dependent cDC1 activation and tumor accumulation of endogenous Tpex and Tdif CD8^+^ T cells, which collectively provide robust and long-term antitumor immunity. Our findings also support the use of targeted lymphodepleting preconditioning regimens that specifically eliminate dispensable or suppressive cell populations, such as Tregs, while sparing host antitumor T cells and cDC1s that may lead to more effective ACTs against solid tumors. In summary, this study demonstrates the multifaceted role of the host immune system in determining the efficacy of ACT. Therefore, a comprehensive understanding of the interplay between the transferred cells and the host immune system is crucial to achieve the full potential of ACT and improve patient outcomes.

## MATERIALS AND METHODS

### Study Design

The overall objective of the study was to test the hypothesis that endogenous CD8^+^ T cells play a key role in ACT-mediated tumor eradication. Using different mouse models, we evaluated the requirement of endogenous T cells for tumor elimination induced by adoptive transfer of *in vitro* activated OTI CD8^+^ T cells. The frequencies and phenotype of endogenous and adoptively transferred tumor-infiltrating T cell populations were analyzed by flow cytometry. Sample size was determined via a priori power analysis based on means and SDs estimated from pilot experiments and previous experience. All mice were randomized before treatment initiation, studies were terminated at a defined end point as indicated or when tumor volume reached 1500 mm^3^, and evaluation of tumor volume was blinded when possible.

### Animals

C57BL/6/J wild-type (CD45.2), B6.129S7-*Rag1^tm1Mom^*/J (RAG1KO), C57BL/6-Tg(TcraTcrb)1100Mjb/J (OT-I), CBy.SJL(B6)-*Ptprc^a^/J* (CD45.1), B6.129S2-Cd207^tm3(DTR/GFP)Mal^/J (Langerin-DTR) mice were purchased from Jackson Laboratories. Mice were kept at the animal facility of Fundación Ciencia & Vida and maintained according to the “Guide to Care and Use of Experimental Animals, Canadian Council on Animal Care”. This study was carried out in accordance with the recommendations of the “Guidelines for the Welfare and use of Animals in cancer research, Committee of the National Cancer Research Institute”. All procedures complied with all relevant ethical regulations for animal research and were approved by the “Comité de Bioética y Bioseguridad” of Fundación Ciencia &Vida. Blinding or randomization strategies were done whenever possible, no animals were excluded from the analysis, and male and female mice were used indistinctly. Mice were allocated randomly in the different experimental procedures.

### Cell lines

Mouse melanoma cell line B16F10 (ATCC CLR-6475) was obtained from American Type Culture Collection. B16F10-OTIx5-ZsGreen (B16F10-OTI) cells were generated by lentiviral transduction of B16F10 cell line with the pLVX-OTIx5-ZsGreen vector encoding the OTI epitope minigene fused to ZsGreen (38). B16F10-OTI cell line was cultured in complete RPMI 1640 (ThermoFisher Scientific, ref 61870-036) media, supplemented with penicillin, streptomycin (ThermoFisher Scientific, ref 15140122), non-essential amino acids (ThermoFisher Scientific, ref11140050), sodium pyruvate (ThermoFisher Scientific, ref 11360070) and 10% of heat-inactivated fetal bovine serum (ThermoFisher Scientific, ref 10437010) in a humidified incubator at 37 °C with 5% CO2. Cell line was routinely tested for mycoplasma contamination.

### Tumor challenge

Mice were injected intradermally in the lower flank with 50μL of PBS containing 1 × 10^6^ of tumor cells. Tumor growth was monitored by measuring perpendicular tumor diameters with calipers. Tumor volume was calculated using the following formula: V= (D x d2)/2 where V is the volume (mm^3^), D is the larger diameter (mm), and d is the smaller diameter (mm). Mice were sacrificed when moribund or when the mean tumor diameter was ≥15mm, according to the approved ethical protocol.

### Adoptive Cell Therapy

To generate OTI-specific CD8^+^ T cells for ACT, splenocytes from OTI mice were cultured in RPMI 1640 supplemented media containing 2 μg/mL of SIINFEKL peptide, 100 UI/mL of recombinant human IL-2 (rhIL-2; Biolegend, ref 589108) and 50 μM of 2-mercaptoethanol (Merck, ref M3148) and expanded daily for 96 hrs. Seven days after tumor challenge, when tumors reached a size of ∼50-200 mm^3^, mice were intravenously injected with 1x10^6^ (optimal) or 0.5x10^6^ (suboptimal) activated OTI CD8^+^ T cells in 100 μL of sterile PBS (ThermoFisher Scienfic, ref 10010023).

### FTY720 treatment, antibody administration, lymphodepleting preconditioning regimen with cyclophosphamide and DC depletion

To block T cell circulation, 25 μg of FTY720 (Sigma-Aldrich, ref SML0700) was injected intraperitoneally every three days starting one day after the tumor challenge. To deplete CD8^+^, one day after the tumor challenge, mice were intraperitoneally injected with three doses of 20 μg of rat monoclonal anti-CD8α antibody (BioXCell, clone YTS169.4, ref BE0117) on consecutive days. Lymphodepleting preconditioning regimen prior to ACT was performed by two intraperitoneal doses of 300 mg/kg cyclophosphamide (Sigma-Aldrich, ref. PHR1404) on day 4 and 6 after tumor challenge. TNF-α blockade was assessed by intraperitoneal injections of 500 μg of anti-TNF-α (Bioxcell, clone XT.11, ref BE0058) every other day starting one day before ACT. To deplete cDC1s, Langerin-DTR mice received 1 μg of diphtheria toxin (Sigma–Aldrich, ref D0564 1MG) by intravenous injection in the tail vein three days after the tumor challenge and continuously maintained doses of 0.35 μg intraperitoneally every 3 days.

### Preparation of tissue cell suspensions

Tumors, inguinal lymph nodes and skin samples were excised, cut in small fragments and mechanically disaggregated. Samples were resuspended in 1 mL RPMI 1640 medium (ThermoFisher Scientific, ref 61870-036) containing 5 mg/mL of collagenase type IV (Gibco, ref 17104019) and 5 μg/mL of DNAse I (AppliChem, ref A3778,0010) and incubated for 60 (tumors) or 30 (lymph nodes and skin) min at 37 °C with shaking. Samples were then resuspended in 1 mL of supplemented of RPMI 1640 medium (ThermoFisher Scientific, ref 61870-036) containing 5 μg/mL of DNAse I (AppliChem, ref A3778,0010) and incubated for 5 min at 4°C. Skin pieces were mechanically disaggregated using microscope slides with ground edges (Sail Brand, ref 7105). Single cell suspensions were obtained using a 70μm cell strainer (BD Falcon, ref 352350). For the analysis of tumor DCs, CD45 magnetic positive selection (MACS, Miltenyi ref 130-052-301) was used to enrich hematopoietic cells.

### Surface, intracellular and intranuclear staining for flow cotometry

Single cell suspensions were incubated for 10 min with the TruStain fcX (Biolegend, clone 93, ref 101320). For surface staining, cell suspensions were incubated with the antibodies for 20 minutes at 4°C followed by two washes with PBS. Cells were then fixed and permeabilized for intracellular and intranuclear staining using the eBioscience FOXP3/transcription factor staining kit (Invitrogen, ref 00-5523-00), followed by intranuclear staining. Monoclonal antibodies specific for mouse molecules were purchased from Biolegend: CD45-FITC (clone 30-F11), CD279/PD1-PE (clone 29f.1a12), CD64-PEDazzle (X54-5/7.1) CD8α -PerCP (clone 53-7.6), PerCP-CD3 (17A2), PerCP-B220 (RA3-6B2), CD45.1-PE/Cy7 (clone A20), CD366/TIM3-PE/Cy7 (clone RMT3-23), PDL1-PE/Cy7 (10F.9G2), CD45.2-APC/Cy7 (clone 104), MHCII-APC/Cy7 (m5/114.15.2), CD3-Brillant Violet 421 (clone 17A2), CD45.1-Brillant Violet 421 (clone A20), CD279/PD1-Brillant Violet 421 (clone 29f.1a12), CD24-Brillant Violet 421 (M1/69), CD366/TIM3-Brillant Violet 605 (RMT3-23), Ly6C-Brillant Violet 605 (HK1.4), CD8-Brillant Violet 650 (53-6.7), XCR1-Brillant Violet 650 (ZET), CD11c-Brillant Violet 711 (N418), CD11b-Brillant Violet 785 (M1/70). BD Biosciences: CD86-PE (cloneGL1). Cell signaling technology: TCF1/TCF7-AF488 (clone C63D9). Miltenyi Biotec: TOX-APC (REA473) and Invitrogen: Granzyme B-APC (clone GB11), Granzyme B-PE-TexasRed (clone GB11). Viability dye was made with ZombieAqua (Biolegend ref 423101). Samples were acquired in a BD FACSCanto II cytometer or BD FACSAria III (BDBioscience), and data were analyzed using FlowJo version 10.8.1 (Tree Star, Inc.).

### Gene set enrichment analysis

To explore whether gene signatures correlate with clinical response in patients under ACT, we performed a Gene Set Enrichment Analysis (GSEA). Normalized gene expression matrix and clinical data were downloaded from the Gene Expression Omnibus (accession number GSE100797) (49). Patients were divided according to RECIST criteria: complete response (CR), partial response (PR), stable disease (SD), and progressive disease (PD). Signatures were obtained and manually curated from previously reported single-cell RNA-seq data and signature databases (11,50,61–64). For enrichment analysis, we used the GSEA software (Broad Institute, v4.2.3), with 1000 permutations, and gene set as permutation type (65) Plot was generated with ggplot2 (v3.4).

### TCGA survival analysis

To determine whether gene signatures correlate with patients’ survival, we analyzed The Cancer Genome Atlas – Skin Cutaneous Melanoma cohort. Expression and clinical data were downloaded from the Xena Browser (UCSC) (Goldman et al. 2020). For each signature, we calculated the mean log_2_(FPKM-uq + 1). Patients with inconsistent expression and clinical data were filtered. Survival analysis was performed with the “survival” package (v3.4). Patients were divided in high and low according to the gene signature optimal cut point determined with the maximally selected rank statistics method. Kaplan-Meier survival curves were drawn with the “survminer” package (v3.2). All analyses were carried out in R environment (v4.1.1). A p<0.05 was considered as statistically significant.

### Statistical analyses

Statistical analyses were performed using Graphpad Prism software (Graphpad Software Inc.). Mann-Whitney unpaired test was performed between relevant groups. Statistical analysis for tumor growth was performed using two-way ANOVA Bonferroni post-hoc test. Error bars in the figures indicate the mean plus SEM. Survival analysis was done by Kaplan-Meier curve with a Log-rank test. Overall p value <0.05 was considered statistically significant; *p≤0.05, **p≤0.01, ***p≤0.001 and ****p≤0.0001.

## Funding

Centro Cientifico y Tecnologico de Excelencia Ciencia & Vida, FB210008 from ANID - Agencia Nacional de Investigación y Desarrollo (AL, VB, HG).

FONDECYT grants 1212070 (AL), 11190897 (VB), 3220741 (AHO), 3220788 (SH), 3240458 (DF), and 3240490 (XL) from ANID - Agencia Nacional de Investigación y Desarrollo.

PhD Scholarship 21180968 (DF) from ANID - Agencia Nacional de Investigación y Desarrollo.

## Author Contributions

Conceptualization: DF, AL

Methodology: DF, FGC, AHO, and FF

Investigation: DF, FGC, AHO, JPV, SH, FA, FF, and XL

Funding acquisition: HG, VB, AL

Project administration: VB, AL

Supervision: FO, VB, AL

Writing – original draft: DF, AL,

Writing – review & editing: DF, FGC, AHO, FO, AL,

## Competing Interests

The authors declare no competing interests.

## Data and materials availability

All data are available in the main text or the supplementary materials.

## SUPLEMENTARY FIGURES

**Supplementary Figure 1.**
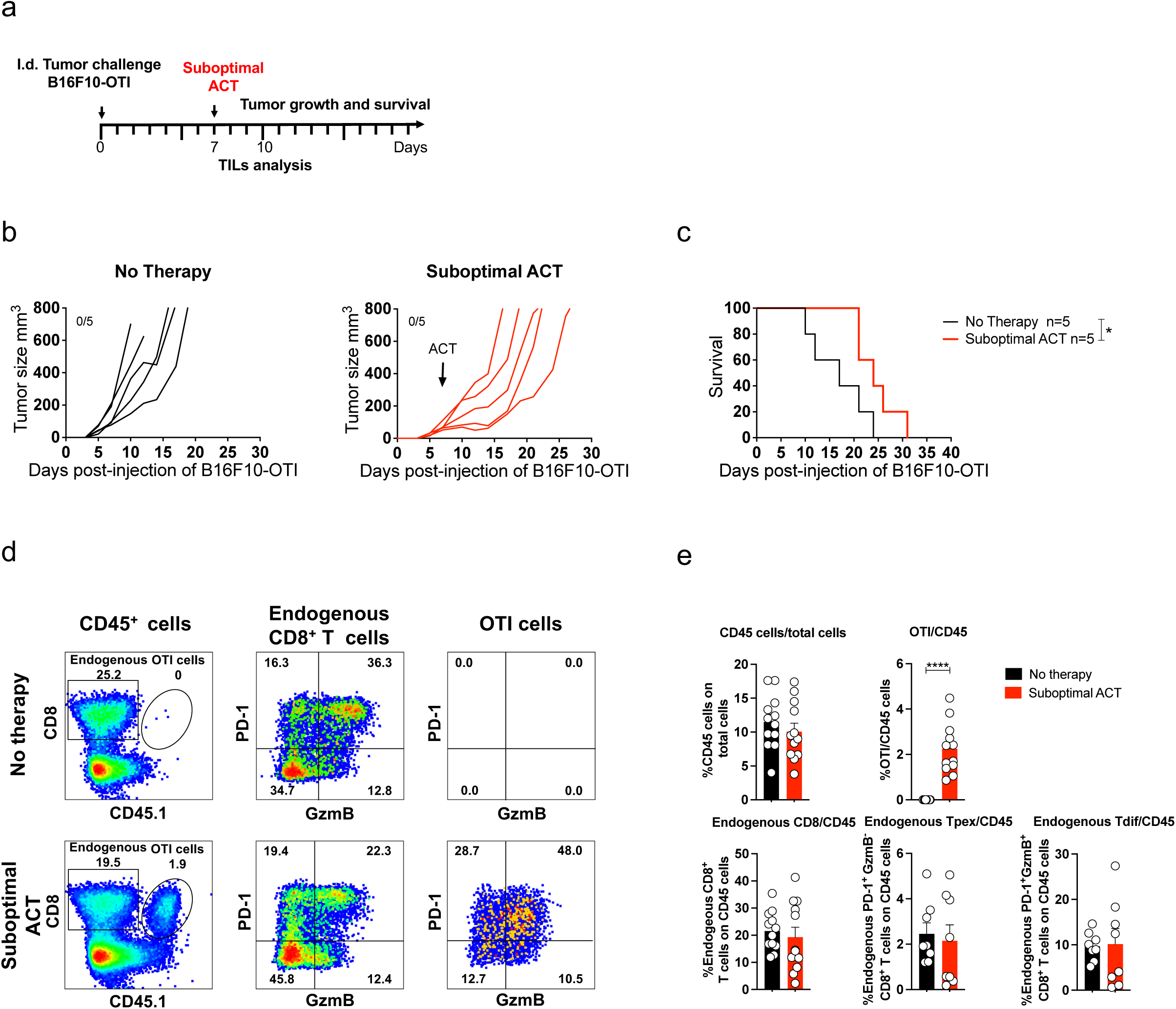
Suboptimal ACT does not promote tumor accumulation of endogenous CD8 T cells and lead to tumor progression. C57BL/6 mice bearing B16F10-OTI tumors received i.v. transfer of 0.5x10^6^ *in vitro* activated OTI CD8^+^ T cells as a Suboptimal ACT. Untreated mice (No therapy) were used as controls. **a** Experimental scheme. **b-c** Individual tumor growth (**b**) and Kaplan-Meier (**c**) curves for each group: No therapy (black curves) and Suboptimal ACT (red curves). **d-e** Tumor infiltrates were analyzed by flow cytometry three days after ACT. **d** Left panels: Representative dot plots displaying the frequencies of endogenous (square, CD45.1^-^) and transferred (ellipse, OTI CD45.1^+^) CD8^+^ T cells in live CD45^+^ cells. Middle panels: Representative dot plots displaying the expression of PD-1 and granzyme B in endogenous CD8^+^ T cells and the frequencies of the different subpopulations defined in each quadrant: PD-1^+^ GzmB^-^ (Tpex), PD-1^+^ GzmB^-^ (Tdif), PD-1^-^ GzmB^-^ and PD-1^-^ GzmB^+^. Right panels: Representative dot plots displaying the expression of PD-1 and granzyme B in transferred OTI CD8^+^ T cells. **e** Quantifications of CD45^+^ cells as a percentage of total live cells, transferred OTI, total endogenous, Tpex (PD-1^+^ GzmB^-^) and Tdif (PD-1^+^ GzmB^+^) CD8^+^ T cells as a percentage of CD45^+^ cells. (**e**) Pooled data from two independent experiments, n = 8 per group. Bars are the mean ± SEM. *p<0.05, **p< 0.01, ***p< 0.001 by Mann-Whitney unpaired.

**Supplementary Figure 2.**
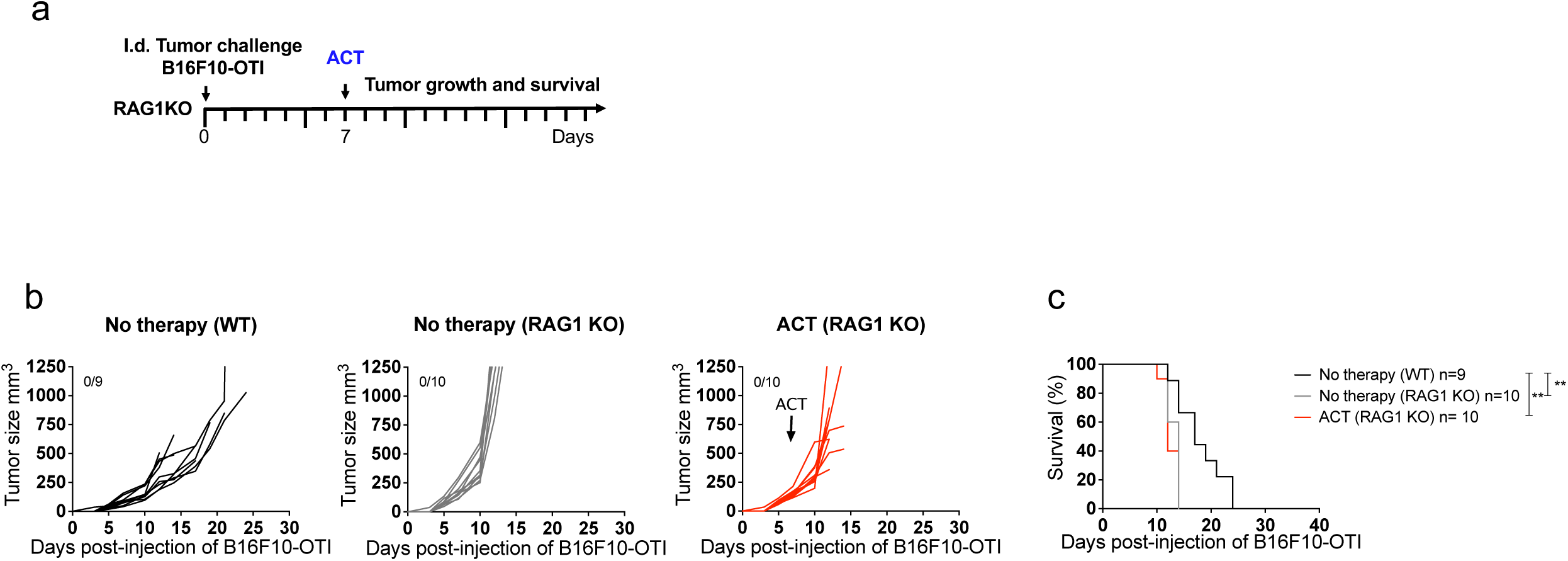
Protection induced by ACT relies on mature/adaptative endogenous immune system. RAG1 KO mice bearing B16F10-OTI tumors received i.v. transfer of 1x10^6^ *in vitro* activated OTI CD8^+^ T cells (ACT). WT untreated mice (No therapy) were used as controls. **a** Experimental timeline. **b-c** Individual tumor growth (**b**) and Kaplan-Meier (**c**) curves for each group: No therapy WT mice (black curves) No therapy RAG1 KO mice (grey curves) and ACT RAG1 KO mice (red curves). (**c**) Results from two independent experiment n=9-10. *p ˂0.05, **p ˂0.01, ***p ≤ 0.001, and ****p ≤ 0.0001 by Log-rank Mantel-Cox test.

**Supplementary Figure 3.**
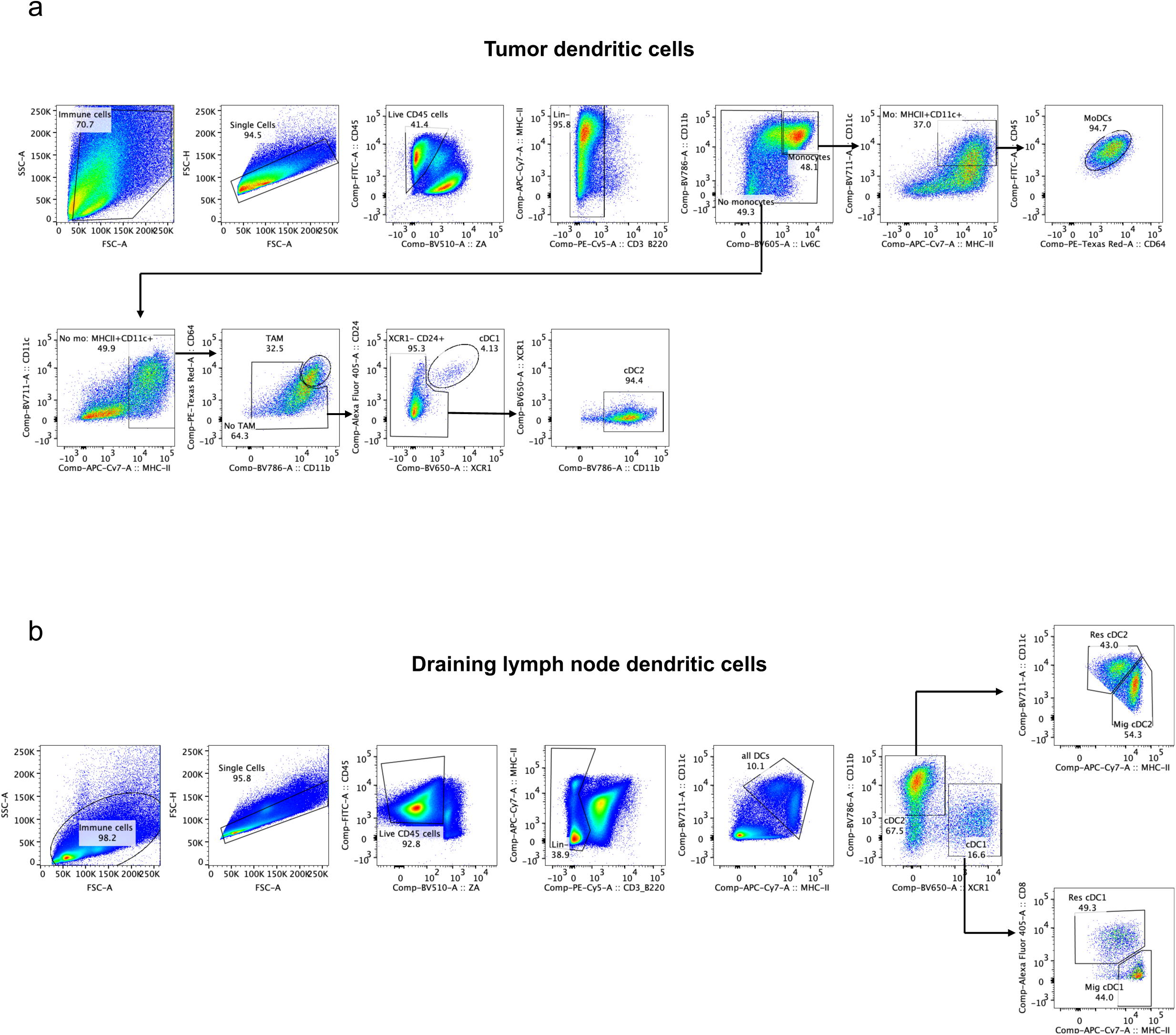
Dendritic cell analysis by flow cytometry in tumor and draining lymph nodes. CD45^+^ cells from tumor and draining lymph node of B16F10-OTI melanoma tumors bearing mice were isolated and staining for flow cytometry analysis. **a** Gating strategy to define tumor-infiltrating type 1 and 2 dendritic cells. **b** Gating strategy to define infiltrating draining lymph node resident and migratory type 1 and 2 dendritic cells.

**Supplementary Figure 4.**
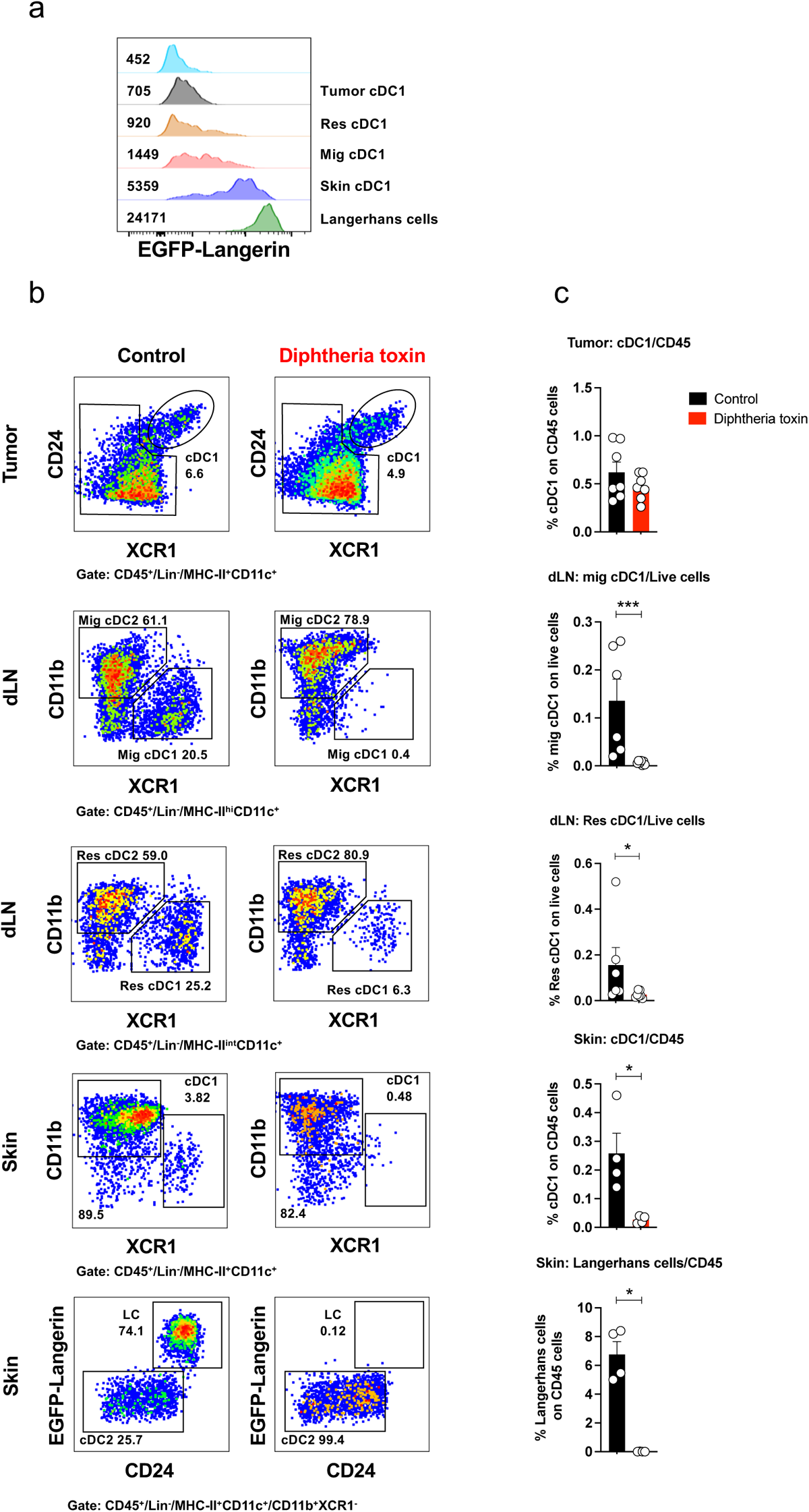
Analysis of dendritic cells from skin, tumor and draining lymph nodes in Langerin-DTR mice. Lang-DTR mice were intradermally challenged with 1x10^6^ B16F10-OTI melanoma tumor cells, then mice received intravenously 1 µg of DTx and three days later tumor, skin and draining lymph node were processed to analyze EGFP-langerin expressing dendritic cells and Langerhans cells. **a** Representative histogram showing the expression of EGFP-Langerin in cDC1 and Langerhans cells across the different analyzed tissues in control untreated DTx B16F10-OTI tumor bearing LANG-DTR mice. **b** Representative pseudocolor dot-plot and frequencies of cDC1 and Langerhans cells in the different tissues in untreated control and DTx treated LANG-DTR mice. **c** Quantification of cDC1 in tumor on CD45^+^ cells, migratory and resident cDC1 in draining lymph node on Live cells, cDC1 and Langerhans cells in skin on CD45^+^ cells. (**c**) Pool of two independent experiments, n = 4-7 per group. Bars are the mean ± SEM. **p ˂0.05, **p ˂0.01, ***p ≤ 0.001, and ****p ≤ 0.0001 by Mann-Whitney unpaired test.

## Notes

### Competing Interest Statement

The authors have declared no competing interest.

## REFERENCES

1. Maier B, Leader AM, Chen ST, Tung N, Chang C, LeBerichel J, et al. A conserved dendritic-cell regulatory program limits antitumour immunity. Nature. 2020 Apr 9;580(7802):257–62.

2. Ruhland MK, Roberts EW, Cai E, Mujal AM, Marchuk K, Beppler C, et al. Visualizing Synaptic Transfer of Tumor Antigens among Dendritic Cells. Cancer Cell. 2020 Jun 8;37(6):786–799.e5.

3. Broz ML, Binnewies M, Boldajipour B, Nelson AE, Pollack JL, Erle DJ, et al. Dissecting the Tumor Myeloid Compartment Reveals Rare Activating Antigen-Presenting Cells Critical for T Cell Immunity. Cancer Cell. 2014 Nov 10;26(5):638–52.

4. Russell JH, Ley TJ. Lymphocyte-mediated cytotoxicity. Vol. 20, Annual Review of Immunology. 2002. p. 323–70.

5. Ariotti S, Hogenbirk MA, Dijkgraaf FE, Visser LL, Hoekstra ME, Song JY, et al. Skin-resident memory CD8+ T cells trigger a state of tissue-wide pathogen alert. Science (1979). 2014 Oct 3;346(6205):101–5.

6. Schenkel JM, Fraser KA, Beura LK, Pauken KE, Vezys V, Masopust D. Resident memory CD8 t cells trigger protective innate and adaptive immune responses. Science (1979). 2014 Oct 3;346(6205):98–101.

7. Khan O, Giles JR, McDonald S, Manne S, Ngiow SF, Patel KP, et al. TOX transcriptionally and epigenetically programs CD8+ T cell exhaustion. Nature [Internet]. 2019 Jul 11 [cited 2021 Apr 14];571(7764):211–8. Available from: https://www.nature.com/articles/s41586-019-1325-x

8. Sekine T, Perez-Potti A, Nguyen S, Gorin JB, Wu VH, Gostick E, et al. TOX is expressed by exhausted and polyfunctional human effector memory CD8+ T cells. Sci Immunol [Internet]. 2020 Jul 1 [cited 2021 Apr 14];5(49). Available from: https://immunology.sciencemag.org/content/5/49/eaba7918

9. Utzschneider DT, Charmoy M, Chennupati V, Pousse L, Ferreira DP, Calderon-Copete S, et al. T Cell Factor 1-Expressing Memory-like CD8+ T Cells Sustain the Immune Response to Chronic Viral Infections. Immunity. 2016;45(2):415–27.

10. Beltra JC, Manne S, Abdel-Hakeem MS, Kurachi M, Giles JR, Chen Z, et al. Developmental Relationships of Four Exhausted CD8+ T Cell Subsets Reveals Underlying Transcriptional and Epigenetic Landscape Control Mechanisms. Immunity. 2020 May;52(5):825–841.e8.

11. Kallies A, Zehn D, Utzschneider DT. Precursor exhausted T cells: key to successful immunotherapy? Vol. 20, Nature Reviews Immunology. Nature Research; 2020. p. 128–36.

12. Siddiqui I, Schaeuble K, Chennupati V, Fuertes Marraco SA, Calderon-Copete S, Pais Ferreira D, et al. Intratumoral Tcf1 + PD-1 + CD8 + T Cells with Stem-like Properties Promote Tumor Control in Response to Vaccination and Checkpoint Blockade Immunotherapy. Immunity. 2019 Jan 15;50(1):195–211.e10.

13. Miller BC, Sen DR, Al Abosy R, Bi K, Virkud Y V., LaFleur MW, et al. Subsets of exhausted CD8+ T cells differentially mediate tumor control and respond to checkpoint blockade. Nat Immunol. 2019 Mar 1;20(3):326–36.

14. Jansen CS, Prokhnevska N, Master VA, Sanda MG, Carlisle JW, Bilen MA, et al. An intra-tumoral niche maintains and differentiates stem-like CD8 T cells. Nature. 2019 Dec 19;576(7787):465–70.

15. Rosenberg SA, Restifo NP. Adoptive cell transfer as personalized immunotherapy for human cancer. Vol. 348, Science. American Association for the Advancement of Science; 2015. p. 62–8.

16. Prickett TD, Crystal JS, Cohen CJ, Pasetto A, Parkhurst MR, Gartner JJ, et al. Durable complete response from metastatic melanoma after transfer of autologous T cells recognizing 10 mutated tumor antigens. Cancer Immunol Res. 2016;4(8):669–78.

17. Stevanovic S, Helman SR, Wunderlich JR, Langhan MM, Doran SL, Kwong MLM, et al. A Phase II Study of Tumor-infiltrating Lymphocyte Therapy for Human Papillomavirus–associated Epithelial Cancers. Clinical Cancer Research. 2019 Mar 1;25(5):1486–93.

18. Scheper W, Kelderman S, Fanchi LF, Linnemann C, Bendle G, de Rooij MAJ, et al. Low and variable tumor reactivity of the intratumoral TCR repertoire in human cancers. Nat Med. 2019 Jan 1;25(1):89–94.

19. Zhang Y, Liu Z, Wei W, Li Y. TCR engineered T cells for solid tumor immunotherapy. Vol. 11, Experimental Hematology and Oncology. BioMed Central Ltd; 2022.

20. Patel U, Abernathy J, Savani BN, Oluwole O, Sengsayadeth S, Dholaria B. CAR T cell therapy in solid tumors: A review of current clinical trials. EJHaem. 2022 Jan;3(S1):24–31.

21. Kochenderfer JN, Somerville RPT, Lu T, Yang JC, Sherry RM, Feldman SA, et al. Long-Duration Complete Remissions of Diffuse Large B Cell Lymphoma after Anti-CD19 Chimeric Antigen Receptor T Cell Therapy. Molecular Therapy. 2017 Oct 4;25(10):2245–53.

22. Porter DL, Levine BL, Kalos M, Bagg A, June CH. Chimeric Antigen Receptor– Modified T Cells in Chronic Lymphoid Leukemia. New England Journal of Medicine. 2011 Aug 25;365(8):725–33.

23. Zhang X, Zhu L, Zhang H, Chen S, Xiao Y. CAR-T Cell Therapy in Hematological Malignancies: Current Opportunities and Challenges. Vol. 13, Frontiers in Immunology. Frontiers Media S.A.; 2022.

24. 24. Morgan RA, Dudley ME, Wunderlich JR, Hughes MS, Yang JC, Sherry RM, et al. Cancer Regression in Patients After Transfer of Genetically Engineered Lymphocytes [Internet]. Available from: www.sciencemag.org

25. Chan JD, Lai J, Slaney CY, Kallies A, Beavis PA, Darcy PK. Cellular networks controlling T cell persistence in adoptive cell therapy. Vol. 21, Nature Reviews Immunology. Nature Research; 2021. p. 769–84.

26. Bechman N, Maher J. Lymphodepletion strategies to potentiate adoptive T-cell immunotherapy–what are we doing; where are we going? Vol. 21, Expert Opinion on Biological Therapy. Taylor and Francis Ltd.; 2021. p. 627–37.

27. Wylie B, Chee J, Forbes CA, Booth M, Stone SR, Buzzai A, et al. Acquired resistance during adoptive cell therapy by transcriptional silencing of immunogenic antigens. Oncoimmunology. 2019;8(8).

28. Khong HT, Restifo NP. Natural selection of tumor variants in the generation of “tumor escape” phenotypes. 2002.

29. O’Donnell JS, Teng MWL, Smyth MJ. Cancer immunoediting and resistance to T cell-based immunotherapy. Vol. 16, Nature Reviews Clinical Oncology. Nature Publishing Group; 2019. p. 151–67.

30. Sotillo E, Barrett DM, Black KL, Bagashev A, Oldridge D, Wu G, et al. Convergence of Acquired Mutations and Alternative Splicing of *CD19* Enables Resistance to CART-19 Immunotherapy. Cancer Discov. 2015 Dec 1;5(12):1282– 95.

31. Braig F, Brandt A, Goebeler M, Tony HP, Kurze AK, Nollau P, et al. Resistance to anti-CD19/CD3 BiTE in acute lymphoblastic leukemia may be mediated by disrupted CD19 membrane trafficking. Blood. 2017 Jan 5;129(1):100–4.

32. Fry TJ, Shah NN, Orentas RJ, Stetler-Stevenson M, Yuan CM, Ramakrishna S, et al. CD22-targeted CAR T cells induce remission in B-ALL that is naive or resistant to CD19-targeted CAR immunotherapy. Nat Med. 2018 Jan 20;24(1):20–8.

33. O’Donnell JS, Teng MWL, Smyth MJ. Cancer immunoediting and resistance to T cell-based immunotherapy. Nat Rev Clin Oncol. 2019 Mar 6;16(3):151–67.

34. Duell J, Leipold AM, Appenzeller S, Fuhr V, Rauert-Wunderlich H, Da Vià MC, et al. Sequential Antigen-loss and Branching Evolution in Lymphoma after CD19- and CD20-Targeted T-cell Redirecting Therapy. Blood Journal. 2023 Nov 17;

35. Jackson HJ, Brentjens RJ. Overcoming antigen escape with CAR T-cell therapy. Cancer Discov. 2015 Dec 1;5(12):1238–40.

36. Lai J, Mardiana S, House IG, Sek K, Henderson MA, Giuffrida L, et al. Adoptive cellular therapy with T cells expressing the dendritic cell growth factor Flt3L drives epitope spreading and antitumor immunity. Nat Immunol. 2020 Aug 1;21(8):914– 26.

37. Walsh SR, Simovic B, Chen L, Bastin D, Nguyen A, Stephenson K, et al. Endogenous T cells prevent tumor immune escape following adoptive T cell therapy. Journal of Clinical Investigation. 2019 Dec 2;129(12):5400–10.

38. Gálvez-Cancino F, López E, Menares E, Díaz X, Flores C, Cáceres P, et al. Vaccination-induced skin-resident memory CD8+ T cells mediate strong protection against cutaneous melanoma. Oncoimmunology [Internet]. 2018 Jul 3 [cited 2021 Apr 17];7(7). Available from: /pmc/articles/PMC5993487/

39. Kaech SM, Cui W. Transcriptional control of effector and memory CD8+ T cell differentiation. Vol. 12, Nature Reviews Immunology. 2012. p. 749–61.

40. Ahn E, Araki K, Hashimoto M, Li W, Riley JL, Cheung J, et al. Role of PD-1 during effector CD8 T cell differentiation. Proceedings of the National Academy of Sciences [Internet]. 2018;115(18). Available from: https://www.pnas.org/content/115/18/4749

41. Chiba K, Matsuyuki H, Maeda Y, Sugahara K. Role of Sphingosine 1-Phosphate Receptor Type 1 in Lymphocyte Egress from Secondary Lymphoid Tissues and Thymus. Vol. 3, Cellular & Molecular Immunology. 2006.

42. Majzner RG, Mackall CL. Tumor antigen escape from car t-cell therapy. Vol. 8, Cancer Discovery. American Association for Cancer Research Inc.; 2018. p. 1219–26.

43. Brunner C, Seiderer J, Schlamp A, Bidlingmaier M, Eigler A, Haimerl W, et al. Enhanced Dendritic Cell Maturation by TNF-α or Cytidine-Phosphate-Guanosine DNA Drives T Cell Activation In Vitro and Therapeutic Anti-Tumor Immune Responses In Vivo. The Journal of Immunology. 2000 Dec 1;165(11):6278–86.

44. Maney NJ, Reynolds G, Krippner-Heidenreich A, Hilkens CMU. Dendritic Cell Maturation and Survival Are Differentially Regulated by TNFR1 and TNFR2. The Journal of Immunology. 2014 Nov 15;193(10):4914–23.

45. Menares E, Gálvez-Cancino F, Cáceres-Morgado P, Ghorani E, López E, Díaz X, et al. Tissue-resident memory CD8+ T cells amplify anti-tumor immunity by triggering antigen spreading through dendritic cells. Nat Commun [Internet]. 2019 Sep 27 [cited 2019 Oct 8];10(1):4401. Available from: http://www.ncbi.nlm.nih.gov/pubmed/31562311

46. Kissenpfennig A, Henri S, Dubois B, Laplace-Builhé C, Perrin P, Romani N, et al. Dynamics and function of langerhans cells in vivo: Dermal dendritic cells colonize lymph node areasdistinct from slower migrating langerhans cells. Immunity. 2005;22(5):643–54.

47. Spranger S, Dai D, Horton B, Gajewski TF. Tumor-Residing Batf3 Dendritic Cells Are Required for Effector T Cell Trafficking and Adoptive T Cell Therapy. Cancer Cell. 2017 May 8;31(5):711–723.e4.

48. Nissani A, Lev-Ari S, Meirson T, Jacoby E, Asher N, Ben-Betzalel G, et al. Comparison of non-myeloablative lymphodepleting preconditioning regimens in patients undergoing adoptive T cell therapy. J Immunother Cancer. 2021 May 14;9(5).

49. Lauss M, Donia M, Harbst K, Andersen R, Mitra S, Rosengren F, et al. Mutational and putative neoantigen load predict clinical benefit of adoptive T cell therapy in melanoma. Nat Commun. 2017 Dec 1;8(1).

50. Jerby-Arnon L, Shah P, Cuoco MS, Rodman C, Su MJ, Melms JC, et al. A Cancer Cell Program Promotes T Cell Exclusion and Resistance to Checkpoint Blockade. Cell. 2018 Nov 1;175(4):984–997.e24.

51. Morotti M, Albukhari A, Alsaadi A, Artibani M, Brenton JD, Curbishley SM, et al. Promises and challenges of adoptive T-cell therapies for solid tumours. Vol. 124, British Journal of Cancer. Springer Nature; 2021. p. 1759–76.

52. Huang J, Khong HT, Dudley ME, El-Gamil M, Li YF, Rosenberg SA, et al. Survival, Persistence, and Progressive Differentiation of Adoptively Transferred Tumor-Reactive T Cells Associated with Tumor Regression. Vol. 28, J Immunother. 2005.

53. Dudley ME, Wunderlich JR, Yang JC, Sherry RM, Topalian SL, Restifo NP, et al. Adoptive cell transfer therapy following non-myeloablative but lymphodepleting chemotherapy for the treatment of patients with refractory metastatic melanoma. Journal of Clinical Oncology. 2005;23(10):2346–57.

54. Klebanoff CA, Khong HT, Antony PA, Palmer DC, Restifo NP. Sinks, suppressors and antigen presenters: How lymphodepletion enhances T cell-mediated tumor immunotherapy. Vol. 26, Trends in Immunology. Elsevier Ltd; 2005. p. 111–7.

55. Boulch M, Cazaux M, Loe-Mie Y, Thibaut R, Corre B, Lemaître F, et al. A cross-talk between CAR T cell subsets and the tumor microenvironment is essential for sustained cytotoxic activity [Internet]. Vol. 6, Sci. Immunol. 2021. Available from: https://www.science.org

56. Nowicki TS, Peters CW, Quiros C, Kidd CK, Kawakami M, Klomhaus AM, et al. Infusion Product TNFa, Th2, and STAT3 Activities Are Associated with Clinical Responses to Transgenic T-cell Receptor Cell Therapy. Cancer Immunol Res. 2023 Dec 10;11(12):1589–97.

57. Schenkel JM, Herbst RH, Canner D, Li A, Hillman M, Shanahan SL, et al. Conventional type I dendritic cells maintain a reservoir of proliferative tumor-antigen specific TCF-1+ CD8+ T cells in tumor-draining lymph nodes. Immunity. 2021 Oct 12;54(10):2338–2353.e6.

58. Connolly KA, Kuchroo M, Venkat A, Khatun A, Wang J, William I, et al. A reservoir of stem-like CD8 + T cells in the tumor-draining lymph node preserves the ongoing antitumor immune response [Internet]. Vol. 6, Sci. Immunol. 2021. Available from: https://www.science.org

59. Li Z, Tuong ZK, Dean I, Willis C, Gaspal F, Fiancette R, et al. In vivo labeling reveals continuous trafficking of TCF-1+ T cells between tumor and lymphoid tissue. Journal of Experimental Medicine. 2022 Jun 6;219(6).

60. Kuhn NF, Lopez A V., Li X, Cai W, Daniyan AF, Brentjens RJ. CD103+ cDC1 and endogenous CD8+ T cells are necessary for improved CD40L-overexpressing CAR T cell antitumor function. Nat Commun. 2020 Dec 1;11(1).

61. Liberzon A, Subramanian A, Pinchback R, Thorvaldsdóttir H, Tamayo P, Mesirov JP. Molecular signatures database (MSigDB) 3.0. Bioinformatics. 2011 Jun;27(12):1739–40.

62. Charoentong P, Finotello F, Angelova M, Mayer C, Efremova M, Rieder D, et al. Pan-cancer Immunogenomic Analyses Reveal Genotype-Immunophenotype Relationships and Predictors of Response to Checkpoint Blockade. Cell Rep. 2017 Jan 3;18(1):248–62.

63. Sade-Feldman M, Yizhak K, Bjorgaard SL, Ray JP, de Boer CG, Jenkins RW, et al. Defining T Cell States Associated with Response to Checkpoint Immunotherapy in Melanoma. Cell. 2018 Nov 1;175(4):998–1013.e20.

64. Villani AC, Satija R, Reynolds G, Sarkizova S, Shekhar K, Fletcher J, et al. Single-cell RNA-seq reveals new types of human blood dendritic cells, monocytes, and progenitors. Science (1979). 2017 Apr 21;356(6335).

65. Subramanian A, Tamayo P, Mootha VK, Mukherjee S, Ebert BL, Gillette MA, et al. Gene set enrichment analysis: A knowledge-based approach for interpreting genome-wide expression profiles [Internet]. 2005. Available from: www.pnas.orgcgidoi10.1073pnas.0506580102

